# A selectable, plasmid-based system to generate CRISPR/Cas9 gene edited and knock-in mosquito cell lines

**DOI:** 10.1101/2020.10.09.333641

**Authors:** Kathryn Rozen-Gagnon, Soon Yi, Eliana Jacobson, Sasha Novack, Charles M. Rice

**Affiliations:** Laboratory of Virology and Infectious Disease, The Rockefeller University, New York, NY 10065, USA

**Author notes:** Corresponding author: Kathryn Rozen-Gagnon.

## Abstract

*Aedes (Ae.) aegypti* and *Ae. albopictus* mosquitoes transmit arthropod-borne diseases around the globe, causing ~700,000 deaths each year. Genetic mutants are valuable tools to interrogate both fundamental vector biology and mosquito host factors important for viral infection. However, very few genetic mutants have been described in mosquitoes in comparison to model organisms. The relative ease of applying CRISPR/Cas9 based gene editing has transformed genome engineering and has rapidly increased the number of available gene mutants in mosquitoes. Yet, *in vivo* studies may not be practical for screening large sets of mutants or possible for laboratories that lack insectaries. Thus, it would be useful to adapt CRISPR/Cas9 systems to common mosquito cell lines. In this study, we generated and characterized a mosquito optimized, plasmid based CRISPR/Cas9 system for use in U4.4 (*Ae. albopictus*) and Aag2 (*Ae. aegypti)* cell lines. We demonstrated highly efficient editing of the *AGO1* locus and isolated knock-down AGO1 cell lines. Further, we used homology-directed repair to establish knock-in Aag2 cell lines with a 3xFLAG-tag at the N-terminus of endogenous *AGO1*. These experimentally verified plasmids are versatile, cost-effective, and efficiently edit immune competent mosquito cell lines that are widely used in arbovirus studies.

## Introduction

Mosquitoes from the genus *Aedes* are worldwide pests and major vectors of arthropod-borne viruses (arboviruses) that cause global human disease^1–3^. Notable members of this genus include *Ae. aegypti,* which transmits a wide variety of arboviruses, and *Ae. albopictus,* which is prevalent in North America and is an emerging vector for certain arboviruses, such as chikungunya virus^2,4,5^. The ability to perform functional genetic studies in mosquitoes and mosquito cells is crucial to our understanding of pro- and anti-viral mosquito host factors and for potential mosquito control strategies^6–8^. Previous genome engineering in mosquitoes has been achieved using transposons^8–11^ and a variety of engineered nucleases such as transcription activator-like effector nucleases (TALENs)^12,13^, zinc-finger nucleases (ZFNs)^14–17^; and homing endonucleases (HEs)^6,18,19^. However, transposon-mediated transgenesis yields imprecise integrations and it can be laborious to engineer nucleases for each target gene. Difficulties modifying mosquito genomes have been compounded by their large and repetitive nature, which makes assembly and annotation a struggle^20–23^. Therefore, despite their importance to human health, loss-of-function mutants in mosquitoes have significantly lagged behind those available in model insects, such as *Drosophila.*

The adaptation of the bacterial type II clustered regularly interspaced short palindromic repeats (CRISPR) and CRISPR-associated sequence 9 (Cas9) immune system for generalized gene editing has revolutionized genome engineering^24–27^. In CRISPR/Cas9 gene editing, the *Streptococcus pyogenes* Cas9 endonuclease is targeted to genomic DNA by complementary guide RNAs, inducing double-stranded breaks (DSBs; for review of CRISPR/Cas9 see^24^). Genomic loci with DSBs stimulate cellular DNA repair machinery that rejoins DSBs by non-homologous end joining (NHEJ). NHEJ disrupts gene function through small insertions or deletions. Alternatively, cellular homology-directed repair (HDR) can be used to correct the gene or insert changes if a homologous donor template is present. The CRISPR/Cas9 system relies on expression of Cas9, a CRISPR RNA (crRNA) that targets genomic DNA adjacent to a protospacer adjacent motif (PAM; NGG motif) and a trans-activating CRISPR RNA (tracr RNA); crRNA and tracrRNA are often provided together as a single guide RNA (sgRNA). Due to its relative ease of adoption and high efficiency, CRISPR/Cas9-mediated gene editing has generated mutants and knock-ins in a wide variety of cells and organisms^28–31^, including *in vivo* in mosquitoes^32–36^ (for review see^37^). CRISPR/Cas9-mediated editing is a significant advance in the toolkit for functional genetic studies in mosquitoes. However, not many laboratories have access to insectaries for *in vivo* experiments, and initial validation of gene function in cells is more practical and cost effective for examining large gene sets. Thus, it is desirable to establish mosquito adapted CRISPR/Cas9 plasmids to generate mutant or knock-in mosquito cell lines; such plasmids have not been reported to-date.

Perhaps due to this lack of mosquito optimized plasmids, there have been relatively few (two) reports of CRISPR/Cas9 edited mosquito cell lines. One study established a clonal cell line (AF5)^38^, which was then used to establish a Dicer-2 defunct AF5 subclone (AF139)^39^. The other generated *Nix* gene loss-of-function and knock-in C6-36 cell lines^33^. However, these reports both relied on *Drosophila* CRISPR/Cas9 plasmids^30^ and contain no information on CRISPR/Cas9 editing efficiency. In the current study, we updated CRISPR/Cas9 plasmids that rely on *Drosophila* promoters^29^ with mosquito promoters for use in mosquito cells. We then applied this system in widely utilized, immune-competent *Ae. aegypti* (Aag2)^40–42^ and *Ae. albopictus* (U4.4)^42,43^ cell lines. Comparing mosquito adapted CRISPR/Cas9 plasmids to previously used *Drosophila* plasmids, we demonstrated increased editing efficiency at the *AGO1* locus. We generated *AGO1-*edited, knock-down U4.4 and Aag2 cell lines, as well as knock-in Aag2 cell lines that contain a 3xFLAG tag at the N-terminus of endogenous *AGO1.* These well-characterized and efficient mosquito optimized CRISPR/Cas9 plasmids will facilitate functional genetic studies in mosquito cell culture systems.

## Results

### Generation and characterization of mosquito optimized CRISPR/Cas9 plasmids

To generate mosquito optimized plasmids for efficient CRISPR/Cas9 gene editing in mosquito cells, we obtained a plasmid used in *Drosophila,* pDCC6^29^. This plasmid relies on two *Drosophila* promoters to express CRISPR/Cas9 components: 1) the RNA Pol III *U6:96Ab*^29,44–48^ *(dme* U6-2) drives transcription of the sgRNA, and 2) the *hsp70Bb*^44,49^ promoter *(dme* phsp70) drives expression of the human codon-optimized *Streptococcus pyogenes* Cas9 (referred to as hSpCas9^26,27,29,50^; Fig. 1a). Because robust hSpCas9 and sgRNA expression is essential for high-efficiency editing, we replaced the *Drosophila* promoters in pDCC6 with appropriate *Ae. aegypti* promoters.

**Figure 1.**
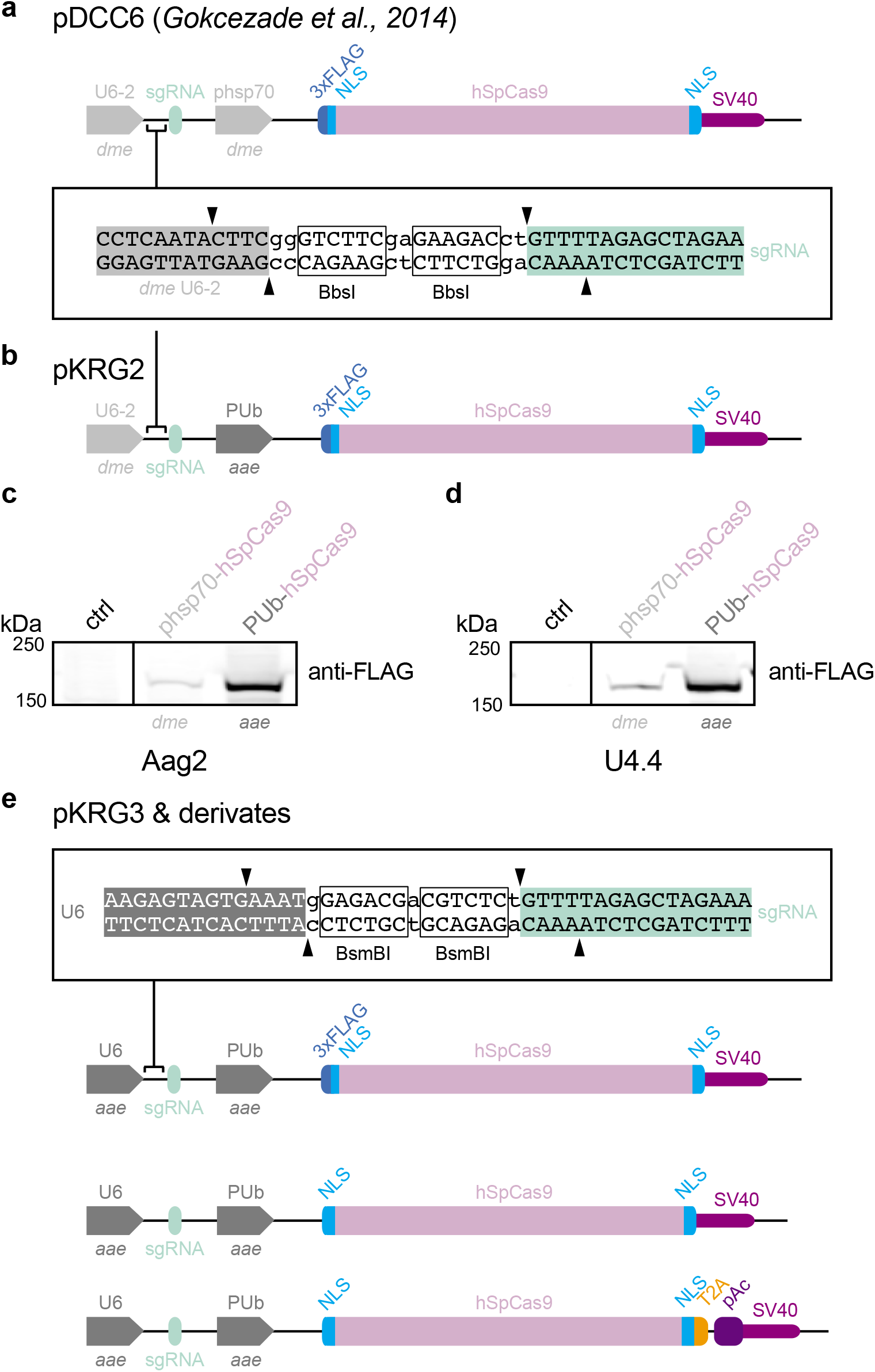
Mosquito optimized CRISPR/Cas9 plasmids. (a) pDCC6 plasmid published in Gokcezade et al., 2014. The *dme* U6-2 promoter drives sgRNA transcription and the *dme* phsp70 promoter drives expression of a 3xFLAG-tagged hSpCas9. Guide RNAs are cloned by *BbsI* digest. (b) pKRG2 plasmid, generated by replacing the *dme* phsp70 promoter with the *aae* PUb promoter. Cloning of guide RNAs as in (a). (c) Immunoblot of Aag2 cells treated with transfection reagent (ctrl) or transfected with pDCC6 (3xFLAG-hSpCas9; expressed from the *dme* pshp70 promoter) or with pKRG2 (3xFLAG-hSpCas9; expressed from the PUb promoter). kDa = kilodaltons. Full-length blots in Supplementary Fig. 3. (d) As in (c) for U4.4 cells. (e) To generate pKRG3 mosquito adapted plasmids, the pKRG2 guide RNA cloning site was redesigned to use *BsmBI* (allowing insertion of the *aae* U6 promoter, which is incompatible with the *BbsI* sgRNA cloning site). The RNA Pol II *aae* U6 promoter was then inserted to express sgRNAs. This plasmid was further modified by removing the N-terminal 3xFLAG from Cas9 and adding a T2A-pAc to allow puromycin selection to yield several options for mosquito optimized CRISPR/Cas9 editing.

We first replaced the *dme* phsp70 with the strong constitutive *Ae. aegypti* polyubiquitin promoter (*aae PUb*^15,32,51^; Fig. 1b). Expression of hSpCas9 in this intermediate plasmid (pKRG2) was then assessed in comparison to the parental pDCC6 plasmid (Fig. 1c,d). We observed increased expression of hSpCas9 in both Aag2 and U4.4 cells using the mosquito PUb promoter. The *dme* phsp70 promoter has long been applied to mosquito cells and, notably, the first stable transformations of mosquito cell lines was performed using this promoter^52^. However, the reported frequency of transformants was quite low, an observation possibly explained by the lower levels of expression we observed from *dme* phsp70 in mosquito cells.

We next updated the U6 promoter to drive sgRNA expression (Fig. 1e). The *Drosophila* U6-2 promoter in the pDCC6 plasmid^29^, which was also used to generate prior CRISPR/Cas9-edited mosquito cells^33,39^, is quite ineffective in mosquito cell lines^53^. We selected the previously described *Ae. aegypti-derived* U6 promoter, *AAEL017774^54^,* which is effective in mosquito cells^53^. These two promoter replacements generate the backbone of the mosquito optimized pKRG3 plasmid (pKRG3-mU6-PUb-3xFLAG-hSpCas9). Because the N-terminal 3xFLAG tag on the hSpCas9 may be undesirable for some applications, we also generated a pKRG3 plasmid with this tag removed (pKRG3-mU6-PUb-hSpCas9).

Finally, we added a selectable marker to pKRG3. Although we optimized transfections (representative transfection of U4.4 shown in Supplementary Fig. 1), no estimates of CRISPR/Cas9-mediated editing efficiency have been reported in mosquito cells. Thus, it may be essential to select transfected cells to increase the likelihood of editing. We generated a plasmid expressing hSpCas9 fused to a selectable marker, a puromycin resistance cassette (pAc). The pAc was inserted after the well-characterized insect virus *Thosea asigna* T2A^55^ ‘2A-like’ ribosome skipping site. This T2A has been applied in *Drosophila* S2 and mosquito (C6/36 and Aag2) cells^30,33,39,56,57^, and was the most efficient skipping site in a comparison of five 2A variations in *Drosophila^58^.* This design ensures co-expression of the hSpCas9 and the pAc selectable marker under one strong, constitutive mosquito promoter (pKRG3-mU6-PUb-hSpCas9-pAc).

To assess the suitability of puromycin selection in Aag2 and U4.4 cells, we performed puromycin kill curves to titrate the optimal concentration (data not shown). Puromycin treatment killed both cell lines efficiently in the absence of pAc expression, and PUb-driven expression of hSpCas9-pAc significantly increased cell viability in the presence of puromycin (Fig. 2a,b). We additionally confirmed efficient T2A skipping in both mosquito cell lines by examining the size of hSpCas9 by immunoblot (Fig. 2c,d). In both cases, the hSpCas9 was processed correctly and we observed a strong band at the expected size (~160 kDa) in both constructs. The slight shift in hSpCas9-pAc is due to additional amino acids in the T2A upstream of the skipped residue, and is consistent with T2A processing in Drosophila^56^. A higher nonspecific band is present in all lanes and does not reflect unprocessed hSpCas9-pAc, which would run at ~185 kDa. Therefore, PUb-Cas9-pAc constructs correctly express and process hSpCas9 and enable rapid selection in U4.4 and Aag2 cells.

**Figure 2.**
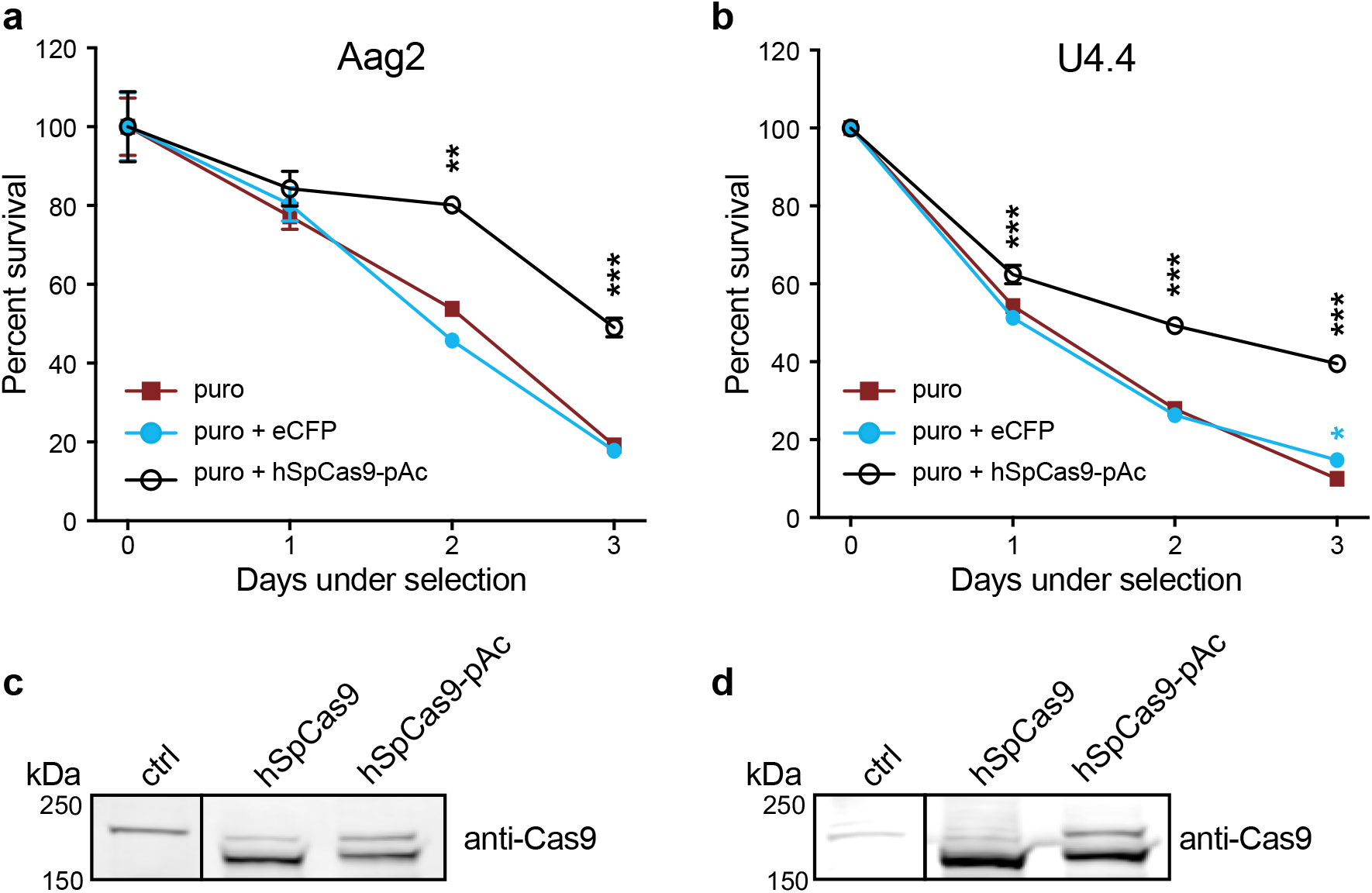
Efficient puromycin selection of hSpCas9-pAc transfected mosquito cells. (a) Aag2 cells were pre-treated with transfection reagent alone (puro), transfected with a control PUb-eCFP (enhanced cyan fluorescent protein) plasmid lacking a puromycin resistance cassette (puro + eCFP), or transfected with PUb-hSpCas9 linked to a puromycin resistance cassette by thes T2A skipping site (hSpCas9-pAc). Day 2 post-transfection, cells were treated with 2.5 ug/mL puro (puromycin). P < 0.0001 (overall ANOVA comparing different transfections); individual groups were compared using the Dunnett’s *post hoc* test compared to the puro control; **P < 0.001, ***P ≤ 0.0001. (b) As in (a) for U4.4 cells treated with 10 ug/mL puro. *P < 0.05, ***P ≤ 0.0001. (c) Immunoblot of Aag2 cells treated with transfection reagent alone (ctrl) or transfected with hSpCas9 or hSpCas9-pAc (driven by PUb promoter). Expected size of hSpCas9 is ~160 kilodaltons (kDa); expected size of unprocessed hSpCas9-pAc is ~185 kDa. Full-length blots in Supplementary Fig. 3. (d) As in (c) for U4.4 cells.

### Efficient CRISPR/Cas9 editing of *AGO1* in U4.4 cells using mosquito-adapted plasmids

As proof-of-principle, we investigated whether the mosquito optimized pKRG3 CRISPR/Cas9 plasmid would enable editing of U4.4 cells at the *AGO1* locus. Guides were designed against the experimentally determined U4.4 genomic sequence near the *AGO1* translational start site by searching for PAM NGG sequences. Because no U4.4 genome has been published, we first verified the starting methionine to target by 5’RACE (this differed from the annotated start site; unpublished results). Three guides were designed in the first exon near the beginning of the coding sequence in order to disrupt *AGO1* (Fig. 3a). Guides were cloned into pKRG3 and these plasmids were transfected singly or in combination into U4.4 cells (Fig. 3b). Post-puromycin selection, we assessed editing efficiency by surveyor assay (Fig. 3c). In this assay, mismatches between annealed wild-type (WT) and edited amplicons leads to cleavage by the surveyor nuclease. Un-transfected WT cells exhibited the expected amplicon at ~350 bp (black arrow; and a smaller, nonspecific band). In contrast, U4.4. cells transfected with pKRG3 plasmids revealed a slightly lower band (red arrow) in addition to the WT band. This indicates amplification of both WT and CRISPR/Cas9 edited *AGO1* loci from pKRG3-transfected cells. We observed that editing was not equally efficient for all single guides, although they were all designed in close proximity. Further, combinatorial transfections with two or three guides performed by far the best^44,47,48^. For a head-to-head comparison, we additionally examined editing using the same hSpCas9 expression and puromycin selection conditions with the *dme* U6-2 promoter (pKRG2) instead of the *aae* U6 promoter (pKRG3). When guides were expressed from the *dme* U6-2 promoter we did not observe any editing, even using all three guides in combination. Therefore, in U4.4 cells the pKRG3 plasmid substantially increases editing efficiency compared to plasmids that rely on *dme* U6-2 for sgRNA expression.

**Figure 3.**
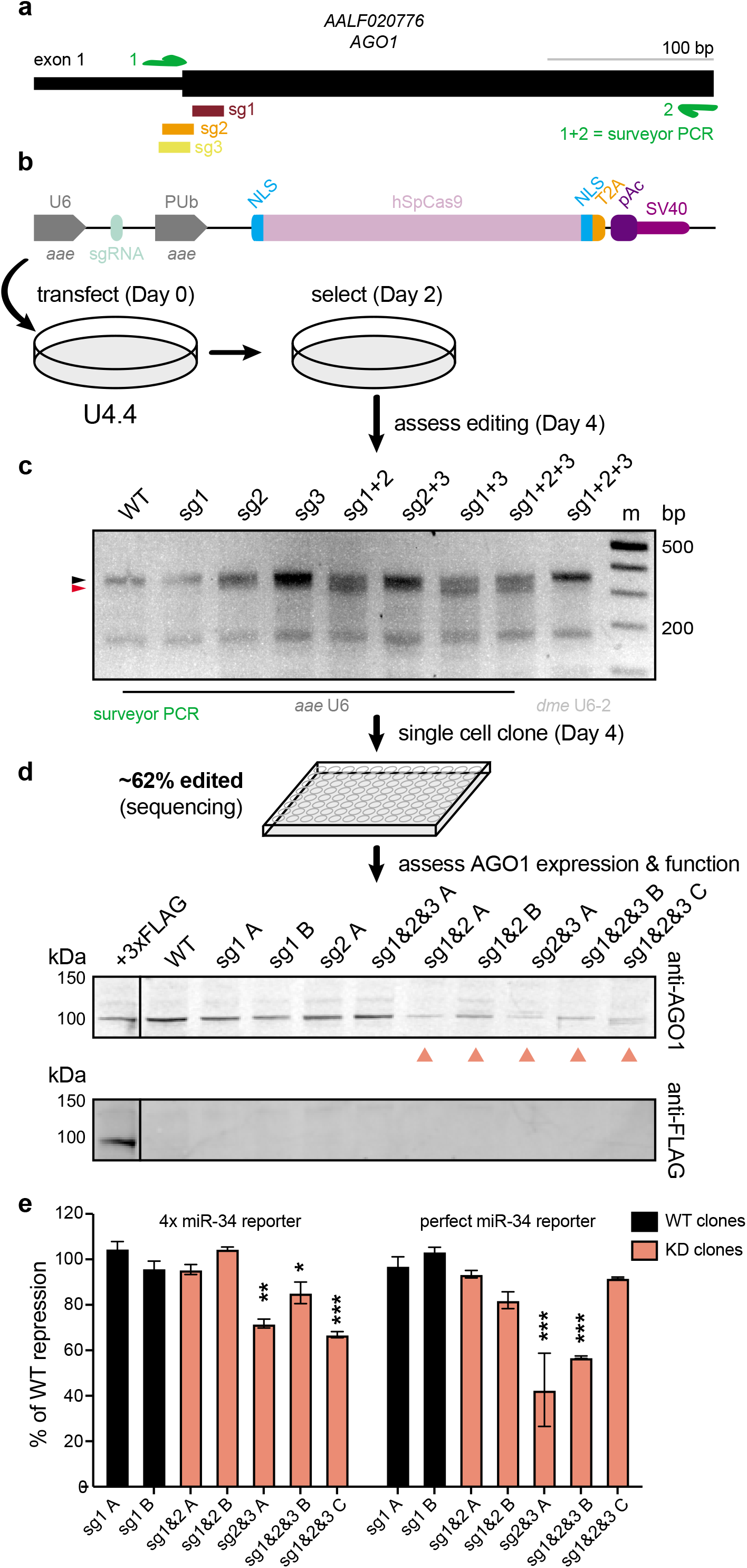
Isolation of AGO1 knock-down U4.4 clonal cell lines. (a) Three sgRNAs were designed near the translation start site for the *AGO1* locus in U4.4 cells. Note that this diagram shown differs from the *Ae. albopictus* annotation (AaloF1.2) and is based on sequencing of U4.4 cells. A PCR was designed for surveyor assay to assess editing efficiency (primers = green arrows). (b) U4.4 *AGO1* sgRNAs were cloned into pKRG3, and U4.4 cells were transfected with these plasmids singly or in combination. Day 2 post-transfection, cells were selected with puromycin until Day 4. (c) Following selection, the surveyor PCR in (a) was used to assess editing by surveyor assay. Expected size of wild-type (WT) amplicon = ~350 base pairs (bp; black arrow); expected size of digested fragments based on sgRNA cleavage sites = ~330 (red arrow) + ~23 (not visible), In comparison to the *aae* U6 promoter, no editing is observed with the same guides using the *dme* U6-2 promoter (both plasmids express hSpCas9 using PUb promoter). m = marker. Full-length gels in Supplementary Fig. 3. (d) Single cells clones were sequenced to assess the percentage of edited clones. Immunoblot of AGO1 (top) showed clones with WT and reduced (salmon arrows) AGO1 protein levels. The ectopically expressed, 3xFLAG-tagged U4.4 AGO1 was detected by the endogenous AGO1 antibody and anti-FLAG antibody (bottom). Full-length blots in Supplementary Fig. 3. (e) Luciferase reporter assay measuring miR-34 mediated repression of 4 repeated ideal miR-34 sites (4x miR-34 reporter) or a perfect miR-34 site (perfect miR-34 reporter). Repression for each clone was measured by normalizing to internal transfection controls (reporter luciferase/control luciferase), then to empty, completely unrepressed reporters in the same clone. The percent (%) of repression compared to WT clones sg1 A and sg1 B is shown. P < 0.0001 (overall ANOVA comparing different transfections); individual groups were compared using the Dunnett’s *post hoc* test compared to the WT clone sg1 A; *p < 0.05, **P < 0.001, ***P ≤ 0.0001.

We next isolated single U4.4 cell clones from this edited population to establish clonal cell lines and assess their potential edits and AGO1 protein level (Fig. 3d). Sequencing of clonal lines showed disruptive edits corresponding to sgRNA cleavage sites (~65% of clones were edited; example alignment shown in Supplementary Fig. 2a). Immunoblot showed decreased protein levels in several of the isolated clones compared to WT levels (Fig. 3d, top). Consistently, we observed that most clones with lower AGO1 levels were isolated from cell populations transfected with combinations of guides. Detection of an ectopically expressed 3xFLAG-tagged U4.4 AGO1 confirmed detection of mosquito AGO1 using the immunoblot antibody (Fig. 3d, bottom). Interestingly, we were unable to isolate a complete knock-out clone despite isolating a variety of different edited lines.

To confirm that reduced AGO1 protein levels corresponded with reduced AGO1 function, we performed a microRNA (miRNA) reporter assay. In mosquitoes, small, host encoded miRNAs direct AGO1 to repress imperfectly complementary transcripts^59^. We assessed whether U4.4 WT or knock-down AGO1 clones could use endogenous miR-34 to repress a *Renilla* luciferase reporter gene containing miR-34 target sequences (Fig. 3e). Several of the knock-down clones, but neither WT clone, exhibited consistently reduced repression. These data indicate that mosquito optimized CRISPR/Cas9 plasmids can be used to edit *AGO1* and to reduce AGO1 protein function in U4.4 cells.

### CRISPR/Cas9 mediates efficient editing and allows knock-in of *AGO1* in Aag2 cells

We next asked whether mosquito adapted plasmids would also allow editing of *Ae. aegypti* Aag2 cells. Additionally, we aimed to generate a knock-in *AGO1* Aag2 cell line by applying CRISPR/Cas9 editing and providing a homology-directed repair (HDR) donor template. To do so, as in U4.4, we designed three guide RNAs near the *AGO1* starting methionine (Fig. 4a). Additionally, we designed an HDR donor template to insert an N-terminal 3xFLAG tag at the endogenous *AGO1* locus (Fig. 4b). Because HDR can be very inefficient, upstream of the 3xFLAG we inserted a red fluorescent protein (RFP) driven by the PUb promoter, flanked by two loxP sites. This allows sorting of RFP positive cells to isolate potentially rare clones with the PUb-RFP-3xFLAG integrated. The PUb-RFP marker can then be excised via expression of Cre recombinase, leaving only a small scar upstream of the 3xFLAG-tagged *AGO1.* This entire cassette was flanked by ~1kb of homology on either terminus^30^.

**Figure 4.**
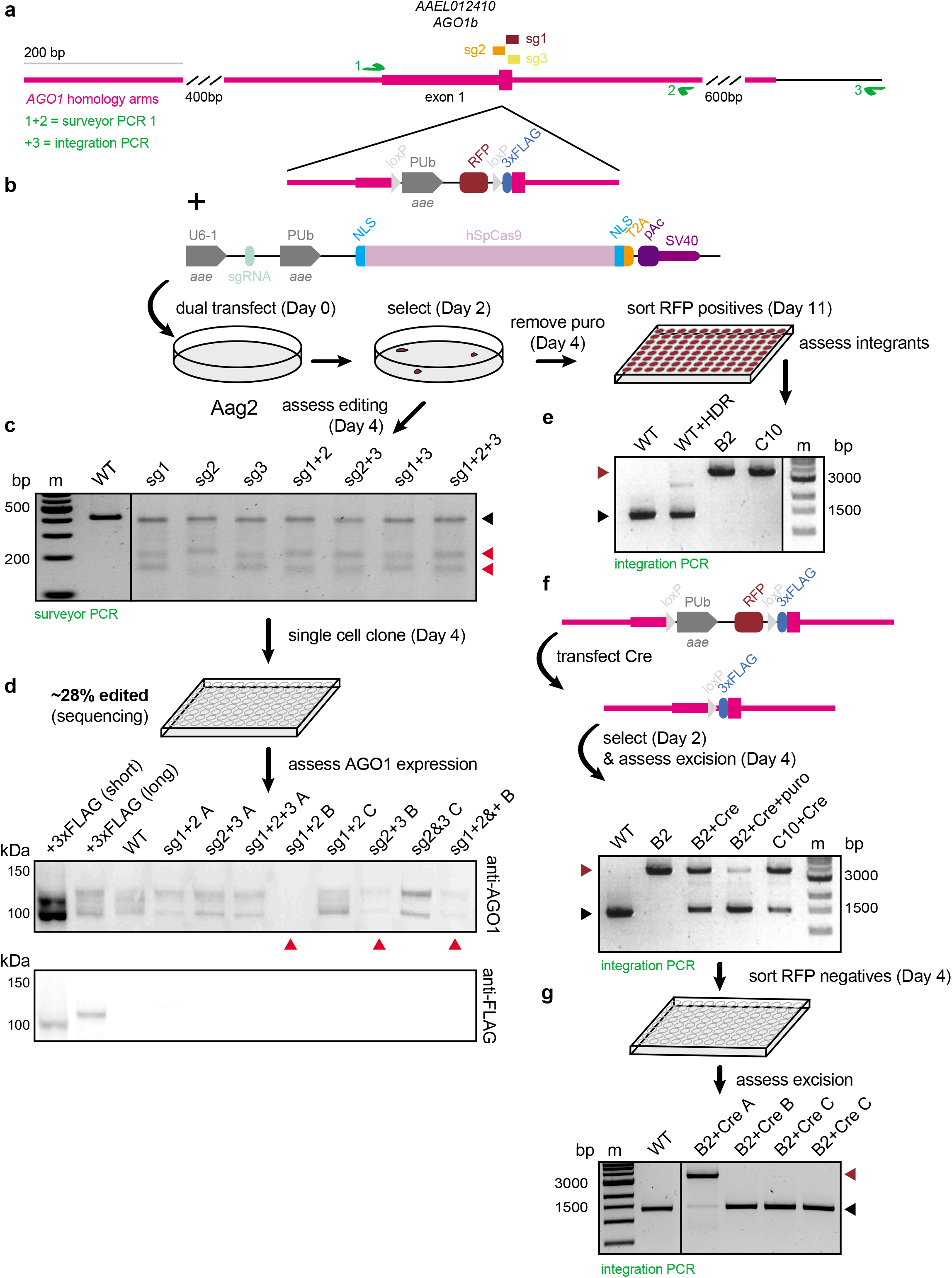
CRISPR/Cas9 mediated editing and knock-in of *AGO1* in Aag2 cells. (a) Three sgRNAs (sg) were designed near the *AGO1* translation start site in Aag2 cells. A PCR was designed for surveyor assay to assess editing efficiency (green arrows 1 and 2). *AGO1* homology arms are indicated; an additional reverse primer was designed outside the homology region (green arrow 3). (b) Homology-directed repair (HDR) donor template design. Upstream of the coding sequence, the PUb promoter expresses red fluorescent protein (RFP); PUb-RFP is flanked by two loxP sites. Aag2 *AGO1* sgRNAs were cloned into pKRG3, and cells were transfected with these plasmids and the HDR donor template. Day 2 post-transfection, cells were selected with puromycin (puro) until Day 4 then screened for editing efficiency. Alternatively, puro selection was removed (Day 4) and cells were cultured until Day 11 and sorted for RFP-positive clones. (c) Following selection, the surveyor PCR in (a) was used to assess editing efficiency by surveyor assay. Expected size of wild-type (WT) amplicon = ~410 base pairs (bp; black arrow); expected size of digested fragments based on sgRNA cleavage sites = ~180 + ~230 (red arrow); m = marker. Full-length gels in Supplementary Fig. 3. (d) Single cell clones were sequenced to determine the percentage of edited clones. Immunoblot of AGO1 (top) showed clones with WT and reduced (red arrows) AGO1 protein levels. The ectopically expressed, 3xFLAG-tagged Aag2 AGO1 short isoform and the 3xFLAG-tagged Aag2 AGO1 long isoform expressed from a stable Aag2 cell line were detected by the endogenous AGO1 antibody and anti-FLAG antibody (bottom). Full-length blots in Supplementary Fig. 3. (e) RFP positive Aag2 cell clones from (b) were screened for HDR mediated integration of the donor template by the integration PCR in (a). Expected size of amplicons from WT clones = ~1500 base pairs (bp; black arrow); expected size of amplicons from clones containing the integrated HDR donor template = ~3,700 bp (dark red arrow). Control WT cells freshly transfected with the HDR donor template (WT+HDR) ensure the donor template is not detected by the integration PCR. B2 and C10 clones were knock-in at the *AGO1* locus. Full-length gels in Supplementary Fig. 3. (f) The PUb-RFP cassette was excised from Aag2 *AGO1* knock-in clones (B2 and C10) by transfection of PUb-driven Cre-T2A-pAc. Transfected cells were selected with puro and the integration PCR in (a) was used to assess excision. WT and transfected B2 cells are shown; expected amplicon sizes as in (e); expected size of Cre excised amplicon = ~1500 base pairs (bp; black arrow). Puro treatment increases the proportion of Cre exicision (B2+Cre+puro). Full-length gels in Supplementary Fig. 3. (g) B2 and C10 Cre excised cell lines were sorted for RFP negative cells and subcloned lines were again screened for homozygous knock-in 3xFLAG-*AGO1* Aag2 cell lines with the integration PCR in (a). Expected amplicon sizes as in (e-f); base pairs = bp. Full-length gels in Supplementary Fig. 3.

Aag2 cells were transfected with pKRG3 plasmids singly or in combination, with the linearized HDR donor template (Fig. 4b). As in U4.4 cells, editing efficiency of bulk Aag2 cells was assessed post-puromycin selection. In contrast to in U4.4 cells, in Aag2 cells we consistently observed high editing efficiency whether guides were expressed singly or in combination (Fig. 4c). Isolation of single cell clones followed by sequencing showed ~28% of isolated clones contained edits (example alignment shown in Supplementary Fig. 2b). Immunoblot showed several clones with decreased and one clone with ablated AGO1 expression (Fig. 4d, top). We were able to detect 3xFLAG-tagged short and long Aag2 AGO1 isoforms by ectopic expression or constitutive expression in stably transformed cell lines, confirming reliable detection of mosquito AGO1 (Fig. 4d, bottom). Unfortunately, the single AGO1 knock-out clone grew poorly and could not be expanded to sufficient cell numbers to establish a knock-out cell line. We were, however, able to establish AGO1 knock-down Aag2 clonal cell lines, as with U4.4.

After transfection and selection, cells were cultured for ~one week, until the input HDR donor template RFP signal diminished. Cells were then sorted to identify clones that were RFP positive due to knock-in of the donor template (Fig. 4b). The efficiency of HDR was very low and only ~0.1% of cells were RFP positives. We screened RFP positive single cell clones by PCR and identified two homozygous clones (B2 and C10) with the correct integration, indicated by a ~3700 bp product (compared to the WT amplicon of ~1500 bp; Fig. 4e). Our primer design outside of the homology arms ensured that this screening PCR only detects the integrated HDR donor template. To excise the loxP-PUb-RFP-loxP cassette, we transfected cells with a plasmid expressing a puromycin-selectable Cre recombinase using the PUb promoter (pKRG4-mPUb-Cre-pAc; Fig. 4f). Upon Cre expression, ~50% of the selectable cassette was excised; this percentage could be increased by selecting for Cre transfected cells with puromycin. To generate the final 3xFLAG-tagged *AGO1* knock-in cell lines, RFP-negatives, which were abundant, were again sorted following Cre transfection and selection (Fig. 4g). PCR of established B2 and C10 subclones indicated homozygous excision. Therefore, CRISPR/Cas9 editing and HDR were applied to commonly used Aag2 cells to knock-in a 3xFLAG tag at the N-terminus of the endogenous *AGO1* locus.

## Discussion

This study provides the first detailed overview and optimization of CRISPR/Cas9 editing in mosquito cells. We generated a set of versatile, selectable CRISPR/Cas9 plasmids, updated for use in mosquito cells. First, by replacing the *Drosophila* hsp70 promoter with the *Ae. aegypti* promoters, we demonstrated increased Cas9 expression in both mosquito cell lines examined. We also verified correct processing of the T2A ribosomal skipping site, enabling constitutive Cas9 and pAc expression from the same PUb promoter (Fig. 2). This strategy was previously employed to generate CRISPR edited Aag2 cells and C6/36 cells, using the same Cas9-T2A-pAc under the *Drosophila* Actin-5c promoter^30,33,39^. We showed that expression of hSpCas9 or Cre fused to T2A-pAc conferred resistance to puromycin and that puromycin treatment increased the desired cell population (Fig. 2, 4). Second, we replaced the *dme* U6-2 promoter with an *aae* promoter for sgRNA expression. Although we saw high levels of editing of the *AGO1* locus using the *aae* U6 promoter, we could not detect any editing using the *dme* U6-2 promoter used in the prior mosquito cell studies^33,39^. This observation is consistent with previous results that the *dme* U6-2 promoter is not very active in mosquito cells^53^. Thus, the mosquito optimized CRISPR plasmids reported here are a significant improvement upon the *Drosophila-based* plasmids previously used in mosquito cells^33,39^.

Interestingly, in contrast to observations in mosquitoes *in vivo*^32^, our results suggest that a single-plasmid CRISPR system works well in mosquito cells. Using the updated mosquito CRISPR/Cas9 plasmids, we obtained ~28-65% edited *AGO1* clones. Co-delivery and expression of both hSpCas9 and sgRNAs from the same, selectable plasmid may increase editing efficiency. Ultimately, high editing efficiencies reduce the labor involved expanding and screening clones, enabling functional interrogation of larger gene sets. Further, selecting bulk edited cells rather than starting with a clonal cell line for editing^39^ allows isolation of multiple WT and knock-out or – down clonal cell lines. Surveying multiple WT and edited clones gives an accurate impression of the phenotypic variation obtained for a given gene mutant from cell to cell. For example, we observed reduced ability to repress miR-34 reporters for the majority of AGO1 knock-down U4.4 clones, but observed variability in the degree of AGO1 impairment. Despite high editing efficiency of the *AGO1* locus in both cell types examined, we were only able to isolate knock-down clones. This is consistent with the impaired growth observed for some miRNA-deficient mammalian cells^60^ and suggests selective pressure against complete *AGO1* knock-outs. Notably, we were also able to use CRISPR/Cas9 plasmids in combination with an HDR donor template to generate Aag2 *AGO1* knock-in cells. Although the efficiency of HDR-mediated integration was low, 0.1% was still sufficiently high to isolate *AGO1* knock-in Aag2 cells. Further, the robust CRISPR/Cas9 system we report will permit future optimization of HDR efficiency in mosquito cells. The ability to knockin large cassettes and excise them allows further flexibility in tagging endogenous proteins for mechanistic studies or generating conditional knock-outs in mosquito cells.

In all, we generated a versatile, effective, single plasmid system for the generation of CRISPR/Cas9 edited mosquito cell lines. The mosquito adapted plasmids we report are a cost-effective tool to screen and investigate functional phenotypes for a large number of gene mutants. This easily customizable set of plasmids can also be updated to encode different Cas9 variants for other applications, such as blocking gene transcription (CRISPR interference) or increasing gene expression (CRISPR activation). It is our hope that this system will facilitate functional genetic studies using widely accessible, immune-competent cell models for major vectors of arboviruses.

## Methods

### Cell lines

Mosquito U4.4 (*Ae. albopictus)* and Aag2^40^ (*Ae. aegypti)* cell lines were kind gifts from Dr. Dennis Brown and from Dr. Maria Carla Saleh, respectively. Cell lines were grown in Leibovitz’s L-15 Medium, no phenol red (Thermo Fisher Scientific), supplemented with 20% (U4.4) or 10% (Aag2) fetal bovine serum (FBS, Hyclone, GE Healthcare), 0.1 mM non-essential amino acids (Thermo Fisher Scientific), and ~0.3g/L tryptose phosphate broth (Sigma-Aldrich) at 28°C, 0% CO_2_.

### Plasmid generation

All CRISPR/Cas9 plasmids were generated from the pDDC6 plasmid, which encodes the human codon-optimized *Streptococcus pyogenes* Cas9 (*hSpCas9*^29,50^; a gift from Peter Duchek (Addgene plasmid #59985; http://n2t.net/addgene:59985; RRID:Addgene_59985). All oligos and gBlocks Gene Fragments were purchased from IDT (see Supplementary Table 1 for all oligo and gBlock sequences). All restriction enzymes, calf intestinal phosphatase (CIP), T4 DNA ligase and Gibson Assembly Master Mix were purchased from NEB, and digests and ligations were performed according to the manufacturer’s protocol. Polymerase chain reactions (PCRs) were performed using Phusion DNA polymerase (NEB) according to manufacturer’s protocols. All transformations were performed using in-house DH5alpha chemically competent cells according to standard protocols. Plasmids were isolated from bacteria using the QIAprep Spin Miniprep Kit (Qiagen) and DNA purifications were performed using QIAquick Gel Extraction and QIAquick PCR Purification Kits (Qiagen), all according to the manufacturer’s protocols. All plasmids were sequence-verified and pKRG3 sequences and the HDR donor template sequence are available in the Supplemental Information.

To replace the *dme* phsp70 promoter (*hsp70Bb*^44,49^) in pDCC6 with the *Ae. aegypti* polyubiquitin promoter (*aae PUb*^15,32,51^), an *AfeI* site was introduced by site-directed mutagenesis using the QuikChange II XL kit (Agilent) and oligos RU-O-22971 and RU-O-22972. The region containing the *Afe1* site was then cloned into a clean pDCC6 background by digestion of the parental pDCC6 and pDCC6-AfeI with flanking sites *SapI/AvrII* followed by ligation, transformation, and DNA isolation. *Aae* Pub with flanking *AfeI/AvrII* sites was amplified using RU-O-22977 and RU-O-22978 and Phusion polymerase (NEB) from the plasmid pSL1180-HR-PUbECFP^15,32^ (a gift from Leslie Vosshall; Addgene plasmid # 47917; http://n2t.net/addgene:47917; RRID:Addgene_47917). The resulting plasmid, pKRG2 (pKRG2-dU6-PUb-3xFLAG-hSpCas9), was sequence verified and contains the *dme* U6-2 promoter (*Drosophila* Pol III promoter *U6:96Ab*^44,45^) and the *aae* PUb promoter driving expression of hSpCas9.

To replace the *dme* U6-2 promoter with the *Aae. aegypti* U6 promoter (*aae* U6; *AAEL017774*^54^), we had to alter the sgRNA cloning sites due to an internal *BbsI* site in the *aae* U6. We designed primers to add an overhang corresponding to the *aae* U6 to a modified sgRNA cloning site that relies on *BsmBI* to the pKRG2 sgRNA tracrRNA (trans-activating CRISPR RNA) scaffold and terminator sequence, with a downstream *Afe*I site (RU-O-22974 and RU-O-22975). The scaffold PCR and a gBlocks Gene Fragment containing the *aae* U6 sequence and an upstream *SacI* site were assembled by Gibson assembly. The assembled DNA was PCR amplified using primers RU-O-22975 and RU-O-22976. The *aae* U6 insert and pKRG2 were digested with *SacI/AfeI,* ligated, and DNA was isolated to obtain pKRG3-mU6-PUb-3xFLAG-hSpCas9, which contains the *aae* U6 promoter driving sgRNA expression and the *aae* PUb promoter driving expression of hSpCas9.

We additionally generated a version of this plasmid with the 3xFLAG at the beginning of the Cas9 removed. To remove the 3xFLAG, we introduced a *NcoI* site by site-directed mutagenesis using oligos RU-O-23100 and RU-O-23101. We then cloned this mutagenized insert into a clean pKRG3 background by restriction enzyme digest with *BglII/XhoI.* The pKRG3-NcoI plasmid was then digested with *NcoI* to remove the 3xFLAG and re-ligated to generate pKRG3-mU6-PUb-hSpCas9. We made another variation with the puromycin resistance cassette (pAc) added. We amplified the end of Cas9, the intervening T2A sequence^56^, and pAc from pAc-sgRNA-Cas9^30^ (a gift from Ji-Long Liu; Addgene plasmid #49330; http://n2t.net/addgene:49330; RRID:Addgene_49330) using primers RU-O-23782 and RU-O-23783. We then digested pKRG3 with *Eag1/BsrGI* and generated pKRG3-mU6-PUb-hSpCas9-pAc by Gibson assembly. For comparative purposes, we also removed the 3xFLAG and added the pAc to hSpCas9 in pKRG3 by the same method, generating pKRG2-dU6-PUb-hSpCas9-pAc.

To generate overexpression plasmids as positive controls for mosquito AGO1 immunoblotting, the pKRG3 plasmid was further modified. Aag2 N-terminal 3xFLAG-tagged short and long AGO1 isoforms and the U4.4 N-terminal 3xFLAG-tagged AGO1 with pKRG3 plasmid overhangs were PCR-amplified from an in-house plasmid containing experimentally validated AGO1 sequences in each cell line (unpublished data). The hSpCas9 sequence was removed from pKRG3-mU6-PUb-hSpCas9-pAc by digestion with *NcoI*/*BsrGI* and AGO1 sequences were inserted by Gibson assembly. Alternatively, to generate an empty pKRG3 plasmid, the ends of the *NcoI/BsrGI*-digested plasmid were filled with T4 DNA polymerase (NEB) according to the manufacturer’s protocol and blunt ends were re-ligated. Finally, the *aae* U6 sequence was removed from pKRG3 Ago-containing or empty plasmids by digestion with *SapI/AfeIl;* the ends were filled with T4 DNA polymerase and blunt ends were ligated. This generated an empty, all-purpose pKRG4-mPUb-pAc plasmid, as well as pKRG4-mPUb-3xFLAG-Aag2-AGO1-short-pAc, pKRG4-mPUb-3xFLAG-Aag2-AGO1-long-pAc, and pKRG4-mPUb-3xFLAG-U44-AGO1-pAc. To express Cre recombinase for excision of the fluorescent reporter between the loxP sites, Cre recombinase was amplified from pME66 (a gift from S. Sarbanes) using primers adding *SacI/AvrII* sites; the Cre insert and pKRG4-mPUb-pAc were digested with *SacI*/*AvrII* to generate pKRG4-mPUb-Cre-pAc.

### CRISPR guide RNA design and cloning

To design CRISPR RNAs (crRNAs) corresponding to *AGO1 (Ae. aegypti* AaegL3 genome assembly, AaegL3.3 annotations, *AAEL012410; Ae. albopictus* AaloF1 assembly, AaloF1.2 annotations, *AALF020776),* we confirmed the genomic sequence around the experimentally determined translational start site in each cell line (unpublished data from 5’RACE and cDNA sequencing). Aag2 and U4.4 cell genomic DNA was isolated using the DNeasy Blood & Tissue Kit (Qiagen) according to the manufacturer’s protocol. Aag2 genomic DNA was amplified using primers RU-O-22776 and RU-O-22777, designed using the *Ae. aegypti* AaegL3 assembly, which has the correct annotated starting methionine. U4.4 genomic DNA was amplified using primers RU-O-22929 and RU-O-22931, designed using the *Ae. albopictus* AaloF1 assembly, where we could only identify a downstream methionine in 5’RACE experiments. Three guide oligos containing the *BsmBI* overhangs in pKRG3 plasmids were designed for each cell line based on protospacer adjacent motif (PAM) NGG sequences in close proximity to the starting methionine (RU-O-23427 to RU-O-23434, *Ae. aegypti*; RU-O-23456 to RU-O-23463, *Ae. albopictus*). The parent pKRG3-mU6-PUb-hSpCas9-pAc plasmid was digested with *BsmBI* and annealed oligos were ligated to generate 6 pKRG3 plasmids, one for each guide, according to protocols from Kistler et al., 2015 and Cornell’s Stem Cell and Transgenic Core Facility (https://transgenics.vertebrategenomics.cornell.edu/genome-editing.html). These co-express the crRNA plus the tracrRNA as a single guide RNA (sgRNA), and hSpCas9.

### Cloning of homology-directed repair (HDR) donor template

The pSL1180-HR-PUbECFP plasmid was used as the backbone for cloning an HDR donor template. A 2kb homology arm fragment around the translational start site of AGO1 in Aag2 cells was amplified from Aag2 genomic DNA using oligos RU-O-24703 and RU-O-24704, to add homology with the pSL1180-HR-PUbECFP plasmid. We ordered a gBlocks Gene Fragment containing an inserted 3xFLAG-tag between the first methionine and the second amino acid of AGO1, with silent mutations to ablate the sgRNA PAM sites. The gBlock extended past *PpuMI/EagI* sites in the homology arm. Next, the homology arm was digested with *PpuMI/EagI* to drop out the central ~160 nt, generating 2 ~1kb homology fragments overlapping the gBlock. pSL1180-HR-PUbECFP was digested with *NotI/EcoRI*, dropping out the PUB-eCFP, and the fragments were assembled to generate the intermediate plasmid pSL1180-HR-Aag2-3xFLAG-AGO1. Next, pSL1180-HR-Aag2-3xFLAG-AGO1 was modified to add the loxp-PUb-RFP-loxP cassette, with overlaps corresponding to the upstream homology arm and downstream 3xFLAG-AGO1 sequence. We generated four PCRs: PCR1) 5’HA-loxP primers = RU-O-25019 and RU-O-25020, template = pSL1180-HR-Aag2-3xFLAG-AGO1 pSL1180-HR-Aag2-3xFLAG-AGO1); PCR2: loxP-PUB-RFP (primers= RU-O-25021 and RU-O-25022, template = pKRG3), PUb-RFP-loxP (primers = RU-O-25023 and RU-O-25024, template= pTRIPZ; Dharmacon), loxP-3xFLAG-3’HA (primers = RU-O-25027 and RU-O-25028, template = pSL1180-HR-Aag2-3xFLAG-AGO1). pSL1180-HR-Aag2-3xFLAG-AGO1 was digested with *KpnI/PpuMI* and the fragments were assembled by Gibson assembly to generate pSL1180-HR-Aag2-loxP-PUb-RFP-loxP-3xFLAG-AGO1.

### General transfections

Aag2 and U4.4 cells were transfected with the appropriate plasmids using Fugene HD Transfection Reagent (Promega) according to the manufacturer’s protocol. Cells were seeded at ~50% confluency and complexes were formed using a ratio of 3:1 transfection reagent to plasmid DNA. In the case of transfection with multiple clones, the total DNA concentration was kept constant and individual plasmids were mixed at equal concentrations.

### Cell survival assays

Cells were seeded at 50% confluency into 96-well plates and transfected with no plasmid, an eCFP control plasmid lacking the pAc (pSL1180-HR-PUbECFP), or a PUb-hSpCas9-pAc plasmid (pKRG2). Day 2 post-transfection, media was replaced with puromycin-containing media (2.5 ug/mL for Aag2 cells and 10ug/mL for U4.4 cells). At Day 0,1,2, and 3 post-puromycin addition, cell survival was analyzed using the CellTiter-Glo Luminescent Cell Viability Assay (Promega) and a FLUOstar Omega Microplate Reader (BMG LABTECH), according to the manufacturer’s instructions. Luciferase values at each day post-puromycin addition were normalized by plasmid paired untreated controls. Survival at each day by plasmid was further normalized to the measured survival for that plasmid at Day 0. Five replicates were collected per condition. Data were analyzed using two-way analysis of variance (ANOVA) with Dunnett’s *post hoc* test, compared to the transfection control without plasmid (Prism 8).

### Generation of stable cell lines

To generate Aag2 cells stably expressing the long AGO1 isoform with an N-terminal 3xFLAG tag, Aag2 cells were transfected with pKRG4-mPUb-3xFLAG-Aag2-AGO1-long-pAc. Day 2 posttransfection, Aag2 cells were treated with a low concentration of puromycin (1 ug/mL) and maintained until cultures recovered (~2 weeks). 3xFLAG-tagged AGO1 expression was confirmed by immunoblot in polyclonal transformants.

### CRISPR/Cas9 transfections

Mosquito cells were transfected with pKRG3-mU6-PUb-hSpCas9-pAc plasmids each containing an AGO1 sgRNA singly, or in combination. Day 2 post transfection, puromycin-containing media was added (2.5 ug/mL, Aag2; 10 ug/mL, U4.4) and cells were selected for an additional 2 days. Post-selection, cell pellets were collected and screened for editing efficiency.

For Aag2 knock-in cells, transfections were performed as above but linearized pSL1180-HR-Aag2-loxP-PUb-RFP-loxP-3xFLAG-AGO1 *(XcmI/AflII)* was co-transfected.

### Surveyor assays

Genomic DNA was isolated from bulk cells transfected with CRISPR/Cas9 plasmids using the epicentre QuickExtract DNA Extraction Solution (Lucigen) according to the manufacturer’s protocol. PCRs were amplified from genomic DNA using primers RU-O-22929 and RU-O-24042 were used for U4.4 cells (full-length PCR product = 353 bp, digested = ~330 bp + ~23 bp); primers RU-O-22776 and RU-O-22777 were used for Aag2 cells (full-length PCR product = 412 bp, digested = ~179 bp + ~231 bp). PCRs were screened for editing efficiency using the Surveyor Mutation Detection Kit (IDT) according to the manufacturer’s instructions, and treated amplicons were visualized on ~1% agarose gels with 100 bp DNA ladder (NEB) and SYBR Gold (Thermo Fisher Scientific).

### Single cell cloning

For bulk CRISPR/Cas9-transfected cells with apparent editing, we isolated single cells to establish clonal edited cell lines. This was done by serial dilution: in brief, cells were resuspended to a concentration of ~7 cells/mL and aliquoted into 96-well plates (~0.7 cells/well) in 50% fresh media, and 50% conditioned media. Cells were expanded and screened by PCR (primers RU-O-22929 and RU-O-24042) and Sanger sequencing (Macrogen).

To isolate knock-in 3xFLAG tagged *AGO1* Aag2 cell lines, cells were resuspended at 2E6/mL in FACS cell sorting buffer (complete L-15 supplemented with 10mM HEPES, 5 mM EDTA and 40ng/mL DAPI). Live, RFP positive cells were sorted for integration of the donor template. Live, RFP negative cells were sorted post Cre transfection and excision. All sorting was performed using a BD FACSAria II sorter (BD Biosciences) with a 100um nozzle and a sheath pressure of ~20 lbf/in^2^ into 96-well plates containing 50% fresh media, and 50% conditioned media. Cells were expanded and genomic DNA was isolated and screened for integration or excision by PCR (primers RU-O-26075 and RU-O-26076); amplicons were visualized on ~1% agarose gels with 1 kb plus DNA ladder (NEB) and SYBR Gold (Thermo Fisher Scientific). WT DNA generates a band of ~1460 bp, DNA with HDR-mediated integration of the donor template generates a band of ~700 bp, and Cre excised DNA generates a band of ~1500 bp). The same PCR was used to Sanger-sequence (Genewiz) B2 and C10 knock-in cell lines to confirm the correct sequence postintegration and post-excision.

### Editing efficiency estimates

The percent of edited clones reported was calculated by sequencing isolated clones (primers RU-O-22776 and RU-O-22777, Aag2; primers RU-O-22930 and RU-O-22931, U4.4). The number of clones that had mixed traces or deletions at sgRNA cleavage sites was calculated over the total number of clones sequenced (Aag2 = 26/93, U4.4 = 8/13).

### SDS-PAGE and immunoblot

Potentially edited clone cell pellets were collected alongside positive control cell pellets. For Aag2 cells, positive controls were Aag2 cells transfected with pKRG4-3xFLAG-Aag2-short or the stable Aag2-3xFLAG-AGO1-long cell line. For U44, cells were transfected with pKRG4-3xFLAG-U44. Pellets were lysed in ice-cold 1X PXL lysis buffer (0.1% SDS, 0.5% sodium deoxycholate, 0.5% NP-40 in 1X PBS) plus protease inhibitor (cOmplete, Mini Protease Inhibitor Cocktail Tablets, EDTA free, Roche) on ice for 10 minutes and clarified by centrifugation at 15,000 rpm for 20 minutes at 4°C. Supernatants were collected and total protein was determined by BCA assay (Pierce BCA Protein Assay Kit, Thermo Scientific) using a FLUOstar Omega Microplate Reader (BMG LABTECH). 10 ug protein/sample was diluted in 1X LDS Sample Buffer and 50 mM DTT (Sigma-Aldrich), then incubated for 10 minutes at 70°C. Samples were loaded on 4-12% Bis-Tris gels and run in 1X MOPS SDS Running Buffer in the Mini Gel Tank according to manufacturer’s Precision Plus Protein Dual Color Standards (Bio-Rad). Following electrophoresis, protein was transferred (Blot Module Set, Thermo Fisher Scientific) onto 0.2 μM nitrocellulose (Amersham Protran Premium, GE Healthcare) according to manufacturer’s protocol. Blots were developed using fluorescent detection (LI-COR) with the following primary antibodies: *anti-Drosophila* AGO1 ab5070 (Abcam), anti-*Ae. aegypti* AGO1 (generated in-house), anti-Cas9 ab204448 (Abcam), and anti-FLAG (monoclonal M2, Sigma-Aldrich). Following development membranes were imaged (Odyssey CLX, LI-COR).

### Luciferase reporter assays

To assay miRNA-mediated silencing in U4.4 knock-down CRISPR clones, reporters were cloned corresponding to 4x miR-34-5p ideal target sites (8mer target followed by 4 mismatches, RU-O- 24800 and RU-O-24801) or one perfect miR-34-5p target site (RU-O-24794 and RU-O-24795). Oligos were annealed and inserted into the *NotI/XhoI-digested* psiCHECK2 plasmid (Promega). 1E5 cells/well were plated in 48-well plates and transfected with empty, perfect, or 4x ideal reporters. Day 2 post-transfection, cells were harvested and analyzed using the Dual-Luciferase Reporter Assay System (Promega) and a FLUOstar Omega Microplate Reader (BMG LABTECH), according to the manufacturer’s instructions. The *Renilla* luciferase (RLuc) signal in each well was normalized to that well’s firefly luciferase (FLuc) signal to measure repression. Then the RLuc/Fluc ratio was normalized to the ratio obtained with the empty reporter in each cell line (which is the unrepressed ratio for that U4.4 clone). To compare normalized ratios to U4.4 clones with WT AGO1 expression, we then divided each clone’s normalized ratio per reporter by the average ratio for that reporter obtained with our two WT clones to obtain a measurement of silencing efficiency. Transfections were performed in triplicate and data were analyzed using twoway analysis of variance (ANOVA) with Dunnett’s *post hoc* test, compared to the WT clone sg1 A (Prism 8).

## Data Availability

The cell lines generated and datasets analyzed during the current study are available from the corresponding author on reasonable request. Plasmid sequences are provided in the Supplementary Information file and pKRG3 and *AGO1* HDR donor plasmids are being deposited to addgene; until the deposit is complete, plasmids are available upon request.

## Acknowledgements

We thank the Rockefeller University Flow Cytometry Resource Center; and M.E. Castillo, S.M. Pecoraro Di Vittorio, A. Norris, S. Shirley, A. O’Connell, and G. Santiago for excellent technical and administrative assistance. We thank the members of the Rice and Vosshall labs and C. McMeniman for invaluable advice, reagents, and discussions. This study was supported by NIH/NIAID grants R01-AI116943 (C.M.R., S.Y, E.J, K.R.G). K.R.G was supported by the Rockefeller University Women and Science Fellowship and a Ruth L. Kirschstein National Research Service Award (NIAID/NIH F32-AI120579).

## Author Contributions

K.R.G and C.M.R. designed the project and experiments. K.R.G., S.Y., E.J., and S.N. performed experiments. K.R.G. performed data analysis and wrote the manuscript with input from all authors.

## Additional Information

The authors declare no competing interests.

## Supplementary Figure Titles and Legends

**Supplementary Figure 1. Efficient transfection of mosquito cells.** (provided in Supplementary Information file)

(a) Representative merged brightfield and CFP image for control (ctrl) U4.4 cells treated with transfection reagent alone.

(b) As in (b), for cells transfected with PUb-eCFP.

**Supplementary Figure 2. *AGO1* sequences of single cell clones isolated after CRISPR/Cas9 transfection.** (provided in Supplementary Information file)

(a) Representative alignment of *AGO1* sequences from established U4.4 single cell clones. Clones were isolated and sequenced post-transfection with pKRG3 CRISPR/Cas9 plasmids containing guides targeting *AGO1.* sgRNA cleavage sites = red arrows; starting methionine = black box.

(b) As in (a), for Aag2 cells.

**Supplementary Figure 3. Full-length blots and gels.** (provided in Supplementary Information file)

(a-j) Full-length blots and gels for all Figures.

**Supplementary Table Titles and Legends**

**Supplementary Table 1. Sequences of all oligos used in this study.** (provided in Supplementary Information file)

HA = homology arm; HDR = homology-directed repair; tracr = trans-activating CRISPR; sgRNA = single-guide RNA. All oligos were ordered in standard desalted format from IDT, unless indicated otherwise.

## References

1. Kraemer, M. U. G. et al. Past and future spread of the arbovirus vectors Aedes aegypti and Aedes albopictus. Nature microbiology 4, 854–863, doi:10.1038/s41564-019-0376-y (2019).

2. Huang, Y.-J. S., Higgs, S. & Vanlandingham, D. L. Emergence and re-emergence of mosquito-borne arboviruses. Current Opinion in Virology 34, 104–109, doi:https://doi.org/10.1016/j.coviro.2019.01.001 (2019).

3. World Health Organization. WHO Fact Sheet: Vector-Borne Diseases, <https://www.who.int/news-room/fact-sheets/detail/vector-borne-diseases> (2020).

4. Schuffenecker, I. et al. Genome microevolution of chikungunya viruses causing the Indian Ocean outbreak. PLoS medicine 3, e263, doi:10.1371/journal.pmed.0030263 (2006).

5. Tsetsarkin, K. A., Vanlandingham, D. L., McGee, C. E. & Higgs, S. A single mutation in chikungunya virus affects vector specificity and epidemic potential. PLoS pathogens 3, e201, doi:10.1371/journal.ppat.0030201 (2007).

6. Windbichler, N. et al. A synthetic homing endonuclease-based gene drive system in the human malaria mosquito. Nature 473, 212–215, doi:10.1038/nature09937 (2011).

7. Hammond, A. M. et al. The creation and selection of mutations resistant to a gene drive over multiple generations in the malaria mosquito. PLoS Genet 13, e1007039, doi:10.1371/journal.pgen.1007039 (2017).

8. Buchman, A. et al. Broad dengue neutralization in mosquitoes expressing an engineered antibody. PLOS Pathogens 16, e1008103, doi:10.1371/journal.ppat.1008103 (2020).

9. Lobo, N. F., Hua-Van, A., Li, X., Nolen, B. M. & Fraser, M. J., Jr. Germ line transformation of the yellow fever mosquito, Aedes aegypti, mediated by transpositional insertion of a piggyBac vector. Insect Mol Biol 11, 133–139, doi:10.1046/j.1365-2583.2002.00317.x (2002).

10. Coates, C. J., Jasinskiene, N., Miyashiro, L. & James, A. A. Mariner transposition and transformation of the yellow fever mosquito, Aedes aegypti. Proc Natl Acad Sci U S A 95, 3748–3751, doi:10.1073/pnas.95.7.3748 (1998).

11. Ivics, Z. et al. Transposon-mediated genome manipulation in vertebrates. Nat Methods 6, 415–422, doi:10.1038/nmeth.1332 (2009).

12. Aryan, A., Myles, K. M. & Adelman, Z. N. Targeted genome editing in Aedes aegypti using TALENs. Methods 69, 38–45, doi:https://doi.org/10.1016/j.ymeth.2014.02.008 (2014).

13. Aryan, A., Anderson, M. A. E., Myles, K. M. & Adelman, Z. N. TALEN-Based Gene Disruption in the Dengue Vector Aedes aegypti. PLOS ONE 8, e60082, doi:10.1371/journal.pone.0060082 (2013).

14. DeGennaro, M. et al. orco mutant mosquitoes lose strong preference for humans and are not repelled by volatile DEET. Nature 498, 487–491, doi:10.1038/nature12206 (2013).

15. McMeniman, Conor J., Corfas, Román A., Matthews, Benjamin J., Ritchie, Scott A. & Vosshall, Leslie B. Multimodal Integration of Carbon Dioxide and Other Sensory Cues Drives Mosquito Attraction to Humans. Cell 156, 1060–1071, doi:10.1016/j.cell.2013.12.044 (2014).

16. Liesch, J., Bellani, L. L. & Vosshall, L. B. Functional and genetic characterization of neuropeptide Y-like receptors in Aedes aegypti. PLoS neglected tropical diseases 7, e2486–e2486, doi:10.1371/journal.pntd.0002486 (2013).

17. Carroll, D. Genome engineering with targetable nucleases. Annu Rev Biochem 83, 409–439, doi:10.1146/annurev-biochem-060713-035418 (2014).

18. Aryan, A., Anderson, M. A. E., Myles, K. M. & Adelman, Z. N. Germline excision of transgenes in Aedes aegypti by homing endonucleases. Sci Rep 3, 1603–1603, doi:10.1038/srep01603 (2013).

19. Stoddard, B. L. Homing endonucleases from mobile group I introns: discovery to genome engineering. Mob DNA 5, 7–7, doi:10.1186/1759-8753-5-7 (2014).

20. Nene, V. et al. Genome sequence of Aedes aegypti, a major arbovirus vector. Science 316, 1718–1723, doi:10.1126/science.1138878 (2007).

21. Timoshevskiy, V. A. et al. Genomic composition and evolution of Aedes aegypti chromosomes revealed by the analysis of physically mapped supercontigs. BMC Biol 12, 27, doi:10.1186/1741-7007-12-27 (2014).

22. Juneja, P. et al. Assembly of the genome of the disease vector Aedes aegypti onto a genetic linkage map allows mapping of genes affecting disease transmission. PLoS Negl Trop Dis 8, e2652, doi:10.1371/journal.pntd.0002652 (2014).

23. Matthews, B. J. et al. Improved reference genome of Aedes aegypti informs arbovirus vector control. Nature 563, 501–507, doi:10.1038/s41586-018-0692-z (2018).

24. Doudna, J. A. & Charpentier, E. Genome editing. The new frontier of genome engineering with CRISPR-Cas9. Science 346, 1258096, doi:10.1126/science.1258096 (2014).

25. Jinek, M. et al. A Programmable Dual-RNA-Guided DNA Endonuclease in Adaptive Bacterial Immunity. Science 337, 816–821, doi:10.1126/science.1225829 (2012).

26. Mali, P. et al. RNA-Guided Human Genome Engineering via Cas9. Science 339, 823–826, doi:10.1126/science.1232033 (2013).

27. Jinek, M. et al. RNA-programmed genome editing in human cells. Elife 2, e00471, doi:10.7554/eLife.00471 (2013).

28. Peng, Y. et al. Making designer mutants in model organisms. Development 141, 4042–4054, doi:10.1242/dev.102186 (2014).

29. Gokcezade, J., Sienski, G. & Duchek, P. Efficient CRISPR/Cas9 Plasmids for Rapid and Versatile Genome Editing in *Drosophila*. G3: Genes\Genomes\Genetics 4, 2279–2282, doi:10.1534/g3.114.014126 (2014).

30. Bassett, A. R., Tibbit, C., Ponting, C. P. & Liu, J.-L. Mutagenesis and homologous recombination in *Drosophila* cell lines using CRISPR/Cas9. Biology Open 3, 42–49, doi:10.1242/bio.20137120 (2014).

31. Hwang, W. Y. et al. Efficient genome editing in zebrafish using a CRISPR-Cas system. Nat Biotechnol 31, 227–229, doi:10.1038/nbt.2501 (2013).

32. Kistler, K. E., Vosshall, L. B. & Matthews, B. J. Genome engineering with CRISPR-Cas9 in the mosquito *Aedes aegypti*. Cell Rep 11, 51–60, doi:10.1016/j.celrep.2015.03.009 (2015).

33. Liu, P. et al. Nix is a male-determining factor in the Asian tiger mosquito *Aedes albopictus*. Insect Biochemistry and Molecular Biology 118, 103311, doi:https://doi.org/10.1016/j.ibmb.2019.103311 (2020).

34. Suzuki, Y. et al. Non-retroviral Endogenous Viral Element Limits Cognate Virus Replication in *Aedes aegypti* Ovaries. Current Biology 30, 3495–3506.e3496, doi:10.1016/j.cub.2020.06.057 (2020).

35. Dong, S. et al. Heritable CRISPR/Cas9-Mediated Genome Editing in the Yellow Fever Mosquito, Aedes aegypti. PLOS ONE 10, e0122353, doi:10.1371/journal.pone.0122353 (2015).

36. Li, M. et al. Development of a confinable gene drive system in the human disease vector Aedes aegypti. eLife 9, e51701, doi:10.7554/eLife.51701 (2020).

37. Hillary, V. E., Ceasar, S. A. & Ignacimuthu, S. in Genome Engineering via CRISPR-Cas9 System (eds Vijai Singh & Pawan K. Dhar) 219–249 (Academic Press, 2020).

38. Fredericks, A. C. et al. Aedes aegypti (Aag2)-derived clonal mosquito cell lines reveal the effects of pre-existing persistent infection with the insect-specific bunyavirus Phasi Charoen-like virus on arbovirus replication. PLOS Neglected Tropical Diseases 13, e0007346, doi:10.1371/journal.pntd.0007346 (2019).

39. Varjak, M. et al. Aedes aegypti Piwi4 Is a Noncanonical PIWI Protein Involved in Antiviral Responses. mSphere 2, doi:10.1128/mSphere.00144-17 (2017).

40. Lan, Q. & Fallon, A. M. Small heat shock proteins distinguish between two mosquito species and confirm identity of their cell lines. The American journal of tropical medicine and hygiene 43, 669–676 (1990).

41. Barletta, A. B. F., Silva, M. C. L. N. & Sorgine, M. H. F. Validation of Aedes aegypti Aag-2 cells as a model for insect immune studies. Parasites & Vectors 5, 148, doi:10.1186/1756-3305-5-148 (2012).

42. Siu, R. W. et al. Antiviral RNA interference responses induced by Semliki Forest virus infection of mosquito cells: characterization, origin, and frequency-dependent functions of virus-derived small interfering RNAs. J Virol 85, 2907–2917, doi:10.1128/JVI.02052-10 (2011).

43. Condreay, L. D. & Brown, D. T. Exclusion of superinfecting homologous virus by Sindbis virus-infected Aedes albopictus (mosquito) cells. Journal of Virology 58, 81–86 (1986).

44. Gratz, S. J. et al. Genome engineering of *Drosophila* with the CRISPR RNA-guided Cas9 nuclease. Genetics 194, 1029–1035, doi:10.1534/genetics.113.152710 (2O13).

45. Wakiyama, M., Matsumoto, T. & Yokoyama, S. *Drosophila* U6 promoter-driven short hairpin RNAs effectively induce RNA interference in Schneider 2 cells. Biochem Biophys Res Commun 331, 1163–1170, doi:10.1016/j.bbrc.2005.03.240 (2005).

46. Xue, Z. et al. Efficient gene knock-out and knock-in with transgenic Cas9 in Drosophila. G3 (Bethesda) 4, 925–929, doi:10.1534/g3.114.010496 (2O14).

47. Ren, X. et al. Optimized gene editing technology for *Drosophila melanogaster* using germ line-specific Cas9. Proceedings of the National Academy of Sciences 110, 19012–19017, doi:10.1073/pnas.1318481110 (2013).

48. Kondo, S. & Ueda, R. Highly Improved Gene Targeting by Germline-Specific Cas9 Expression in *Drosophila*. Genetics 195, 715–721, doi:10.1534/genetics.113.156737 (2013).

49. Loukeris, T. G. Use of the Minos transposable element from Drosophila hydei as a gene transfer vector for Diptera., University of Crete, Greece, (1996).

50. Cong, L. et al. Multiplex Genome Engineering Using CRISPR/Cas Systems. Science 339, 819, doi:10.1126/science.1231143 (2013).

51. Anderson, M. A. E., Gross, T. L., Myles, K. M. & Adelman, Z. N. Validation of novel promoter sequences derived from two endogenous ubiquitin genes in transgenic Aedes aegypti. Insect molecular biology 19, 441–449, doi:10.1111/j.1365-2583.2010.01005.x (2010).

52. Lycett, G. J. & Crampton, J. M. Stable transformation of mosquito cell lines using a *hsp70::neo* fusion gene. Gene 136, 129–136, doi:https://doi.org/10.1016/0378-1119(93)90456-D (1993).

53. Anderson, M. A. E. et al. Expanding the CRISPR Toolbox in Culicine Mosquitoes: *In Vitro* Validation of Pol III Promoters. ACS Synthetic Biology 9, 678–681, doi:10.1021/acssynbio.9b00436 (2020).

54. Konet, D. S. et al. Short-hairpin RNA expressed from polymerase III promoters mediates RNA interference in mosquito cells. Insect Mol Biol 16, 199–206, doi:10.1111/j.1365-2583.2006.00714.x (2007).

55. Donnelly, M. L. L. et al. The ‘cleavage’ activities of foot-and-mouth disease virus 2A site-directed mutants and naturally occurring ‘2A-like’ sequences. Journal of General Virology 82, 1027–1041, doi:https://doi.org/10.1099/0022-1317-82-5-1027 (2001).

56. González, M. et al. Generation of stable *Drosophila* cell lines using multicistronic vectors. Sci Rep 1, 75–75, doi:10.1038/srep00075 (2011).

57. Varjak, M. et al. Characterization of the Zika virus induced small RNA response in *Aedes aegypti* cells. PLOS Neglected Tropical Diseases 11, e0006010, doi:10.1371/journal.pntd.0006010 (2017).

58. Diao, F. & White, B. H. A Novel Approach for Directing Transgene Expression in *Drosophila*: T2A-Gal4 In-Frame Fusion. Genetics 190, 1139–1144, doi:10.1534/genetics.111.136291 (2O12).

59. Bartel, D. P. Metazoan MicroRNAs. Cell 173, 20–51, doi:10.1016/j.cell.2018.03.006 (2018).

60. Luna, Joseph M. et al. Hepatitis C Virus RNA Functionally Sequesters miR-122. Cell 160, 1099–1110, doi:https://doi.org/10.1016/j.cell.2015.02.025 (2015).

