## Supplementary Information for "A selectable, plasmid-based system to generate CRISPR/Cas9 gene edited and knock-in mosquito cell lines"

#### Supplementary Figures

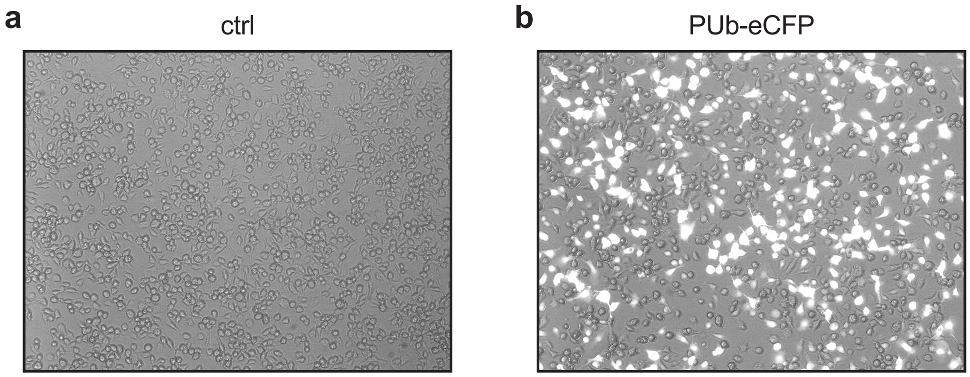


**Supplementary Figure 1. Efficient transfection of mosquito cells.**

(a) Representative merged brightfield and cyan fluorescent protein (CFP) image for control (ctrl) U4.4 cells treated with transfection reagent alone.

(b) As in (a), for cells transfected with PUb-eCFP.

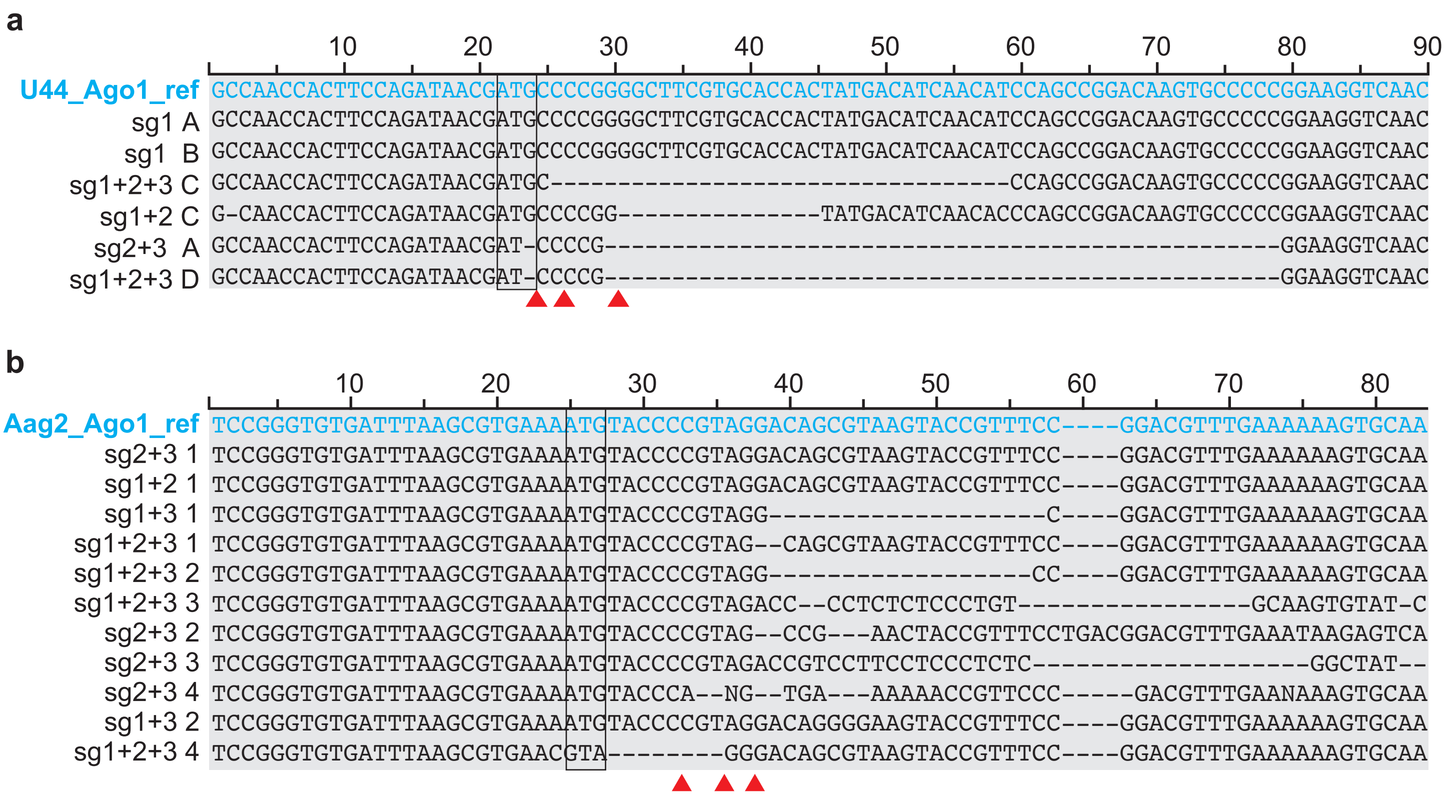


**Supplementary Figure 2. *AGO1* sequences of single cell clones isolated after CRISPR/Cas9 transfection.**

(a) Representative alignment of *AGO1* sequences from established U4.4 single cell clones. Clones were isolated and sequenced post-transfection with pKRG3 CRISPR/Cas9 plasmids containing guides targeting *AGO1*. sgRNA cleavage sites = red arrows; starting methionine = black box.

(b) As in (a), for Aag2 cells.

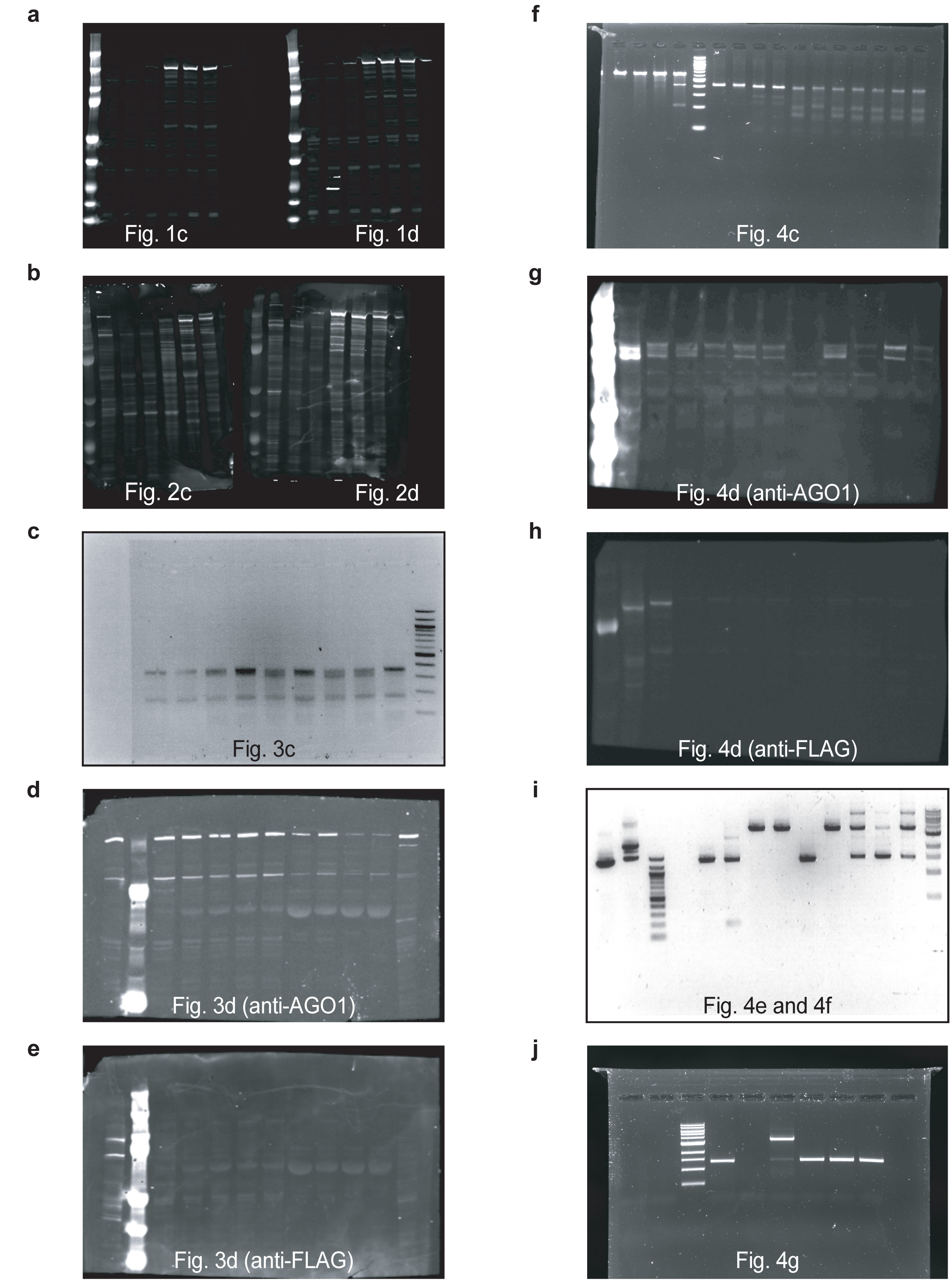


**Supplementary Figure 3. Full-length blots and gels.**

(a-j) Full-length blots and gels for all Figures.

#### Supplementary Tables

##### Supplementary Table 1. Sequences of all oligos used in this study.

| Oligo ID | Sequence | Purpose |
|---|---|---|
| *pKRG cloning* | | |
| RU-O-22971 | CTGCAGAATTGGCGCAAGCGCTAAAAACGGACT | introduce *AfeI* pDCC6 forward; PAGE-purified |
| RU-O-22972 | AGTCCGTTTTTAGCGCTTGCGCCAATTCTGCAG | introduce *AfeI* pDCC6 reverse; PAGE-purified |
| RU-O-22977 | GACAGCGCTTGCGCCAATTCTGCAGACAAATGGCTATCTTTACATGTAGCTTGTGC<br>ATTG | PUb promoter PCR forward |
| RU-O-22978 | GTCCCTAGGTGTATACCTCCGGAAGCGCGCACTCGAGATTCGAACAAGCTTATCGA<br>GCTTGGTGTTGAAATCTCTGTTGAGC | PUb promoter PCR reverse |
| RU-O-22974 | GACGAAGACTATATAAGAGCAGAGGCAAGAGTAGTGAAATGGAGACGACGTCTCTG<br>TTTTAGAGCTAGAAATAGC | tracr RNA scaffold PCR forward |
| RU-O-22975 | GCAGAATTGGCGCAAGCGCTGTC | tracr RNA scaffold & assemble *aae* U6/scaffold PCR reverse |
| RU-O-22976 | GACGAAGAGCGCCCAATACGCAA | assemble aae U6/scaffold PCR forward |
| gBLOCK *aae* U6 | GAAGAGCGCCCAATACGCAAACCGCCTCTCCCCGCGGGTTGGCCGATTCATTAATG<br>CAGGCAACTCGTGAAAGGTAGGCGGATCAGCGAATGAAATCGCCCATCGAGTTGAT<br>ACGTCCATCCATCGCTAGAACCGCGTTCGCTGTAGAAGACTATATAAGAGCAGAGG<br>CAAGAGTAGTGAAATGGAGACG | U6 geneblock |
| RU-O-23101 | GATATTGATTACAAAGACGATGACGATACCATGGCCCCAAAGAAG | introduce *NcoI* remove 3xFLAG forward; PAGE-purified |
| RU-O-23100 | CTTCTTTGGGGCCATGGTATCGTCATCGTCTTTGTAATCAATATC | introduce *NcoI* remove 3xFLAG reverse; PAGE-purified |
| RU-O-25485 | GACCCTAGGATGGGCCCAAAGAAG | Cre into pKRG4 forward |
| RU-O-25486 | CGGTAGAGCTCATCGCCATCTTCCAG | Cre into pKRG4 reverse |
| *sgRNA oligos* | | |
| RU-O-23427 | AAATGGTACTTACGCTGTCCTACG | pKRG Aag2 Ago1 sgRNA 1 forward |
| RU-O-23428 | AAACCGTAGGACAGCGTAAGTACC | pKRG Aag2 Ago1 sgRNA 1 reverse |
| RU-O-23430 | AAATGAGCGTGAAAATGTACCCCGT | pKRG Aag2 Ago1 sgRNA 2 forward |
| RU-O-23431 | AAACACGGGGTACATTTTCACGCTC | pKRG Aag2 Ago1 sgRNA 2 reverse |
| RU-O-23433 | AAATGACGGTACTTACGCTGTCCTA | pKRG Aag2 Ago1 sgRNA 3 forward |
| RU-O-23434 | AAACTAGGACAGCGTAAGTACCGTC | pKRG Aag2 Ago1 sgRNA 3 reverse |
| RU-O-23456 | AAATGTAGTGGTGCACGAAGCCCCG | pKRG U4.4 Ago1 sgRNA 1 forward |
| RU-O-23457 | AAACCGGGGCTTCGTGCACCACTAC | pKRG U4.4 Ago1 sgRNA 1 reverse |
| RU-O-23459 | AAATGTTCCAGATAACGATGCCCCG | pKRG U4.4 Ago1 sgRNA 2 forward |
| RU-O-23460 | AAACCGGGGCATCGTTATCTGGAAC | pKRG U4.4 Ago1 sgRNA 2 reverse |
| RU-O-23462 | AAATGACTTCCAGATAACGATGCCC | pKRG U4.4 Ago1 sgRNA 3 forward |
| RU-O-23463 | AAACGGGCATCGTTATCTGGAAGTC | pKRG U4.4 Ago1 sgRNA 3 reverse |
| *HDR donor template oligos* | | |
| RU-O-24703 | GACCATGATTACGAATTCGACATGTAGGACATGTGGGG | Aag2 HA PCR forward |
| RU-O-24704 | CCAGCTGCAGGCGGCCGCCACCGGATTTGCTTTTCGTC | Aag2 HA PCR reverse |
| gBLOCK Aag2 HDR donor template | CGAAAAGTGCAAAAATTCGGCCGTAATTAGTTCGCTTCCGTTTTTCCGAGTGCCTC<br>CGATCGTTAGCGGACCGTTCCGGGTGTGATTTAAGCGTGAAAATGGATTACAAGGA<br>TCACGATGGAGATTACAAGGATCACGATATCGATTACAAGGATGATGATAAGTATC<br>CGGTGGGTCAACGTAAGTACCGTTTCCGGACGTTTGAAAAAGTGCAATTCACAAG<br>AGAAGAAAAAAAAAGTGTTGACGAGGGGACCTTCCTTCCTCCCTC | |
| RU-O-25019 | GGAGCGTCCGTAAAATGAAA | *Kpn1* HA overlap forward |
| RU-O-25020 | ATAACTTCGTATAGCATACATTATACGAAGTTATTTTCACGCTTAAATC | 5'HA add loxP reverse |
| RU-O-25021 | GCTATACGAAGTTATTATCTTTACATGTAG | loxP-PUb forward |
| RU-O-25022 | GATCAGCTCGCTCATGCGGCCGCGGTAGAGCTCCGAATTCAC | pUB-RFP reverse |
| RU-O-25023 | ATGAGCGAGCTGATC | RFP forward |
| RU-O-25024 | CATTATACGAAGTTATTAATTAATTATCTGTGCCCCAG | RFP-loxP reverse |
| RU-O-25027 | ATAACTTCGTATAATGTATGCTATACGAAGTTATATGGATTACAAGGATC | loxP-HA forward |
| RU-O-25028 | AGAGACGAGAGGGAGGAAGG | HA *PpuMI* overlap reverse |
| *Sequencing, surveyor & integration PCRs* | | |
| RU-O-26075 | ACTTGTCTCACTCGCCATCATACG | Aag2 HDR PCR F |
| RU-O-26076 | CGATCAGATGCGCACGAAAACT | Aag2 HDR PCR R |
| RU-O-22776 | ACTTGTCTCACTCGCCATCA | Aag2 surveyor/sequencing forward |
| RU-O-22777 | GTCCGTACACAGGAAAAGCC | Aag2 surveyor/sequencing reverse |
| RU-O-22929 | GCCAACCACTTCCAGATAACG | U4.4 surveyor forward |
| RU-O-24042 | AGGATGGCATCGTACGGAAT | U4.4 surveyor reverse |
| RU-O-22930 | TCTACCGGTGTTCACCTGTC | U4.4 sequencing forward |
| RU-O-22931 | TGATCTCCCGGTTGACCTTC | U4.4 sequencing reverse |
| *Reporter cloning oligos* | | |
| RU-O-24800 | TCGAGCAACCAGCTAGGCCACTGCCACCGGCAACCAGCTAGGCCACTGCCACCGGC<br>AACCAGCTAGGCCACTGCCACCGGCAACCAGCTAGGCCACTGCCAGC | Aag2 miR-34-5p 4x ideal psiCHECK2 forward |
| RU-O-24801 | GGCCGCTGGCAGTGGCCTAGCTGGTTGCCGGTGGCAGTGGCCTAGCTGGTTGCCGG<br>TGGCAGTGGCCTAGCTGGTTGCCGGTGGCAGTGGCCTAGCTGGTTGC | Aag2 miR-34-5p 4x ideal psiCHECK2 reverse |
| RU-O-24794 | TCGAGCAACCAGCTAACCACACTGCCAGC | Aag2 miR-34-5p perfect psiCHECK2 forward |
| RU-O-24795 | GGCCGCTGGCAGTGTGGTTAGCTGGTTGC | Aag2 miR-34-5p perfect psiCHECK2 reverse |

HA = homology arm; HDR = homology-directed repair; tracr = trans-activating CRISPR; sgRNA = single-guide RNA.
All oligos were ordered in standard desalted format from IDT, unless indicated otherwise.

**Plasmid Sequences**

pKRG3-mU6-PUb-3xFLAG-hSpCas9

*aae* U6
*BbsI* sites for sgRNA cloning
sgRNA tracrRNA and U6 terminator
*aae* PUb promoter
3xFLAG
NLS
hSpCas9

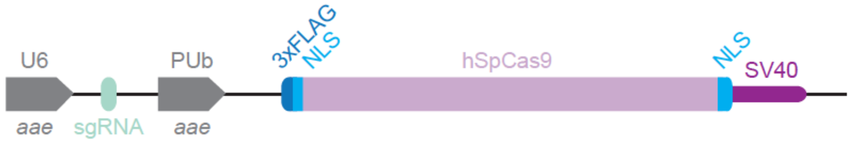


GCGCCCAATACGCAAACCGCCTCTCCCCGCGGGTTGGCCGATTCATTAATGCAGGCAACTCGTGAAAGGTAGGCGGATCAGCGAATGAA
ATCGCCCATCGAGTTGATACGTCCATCCATCGCTAGAACCGCGTTCGCTGTAGAAGACTATATAAGAGCAGAGGCAAGAGTAGTGAAAT
gGAGACGaCGTCTCtGTTTTAGAGCTAGAAATAGCAAGTTAAAATAAGGCTAGTCCGTTATCAACTTGAAAAAGTGGCACCGAGTCGGT
GCTTTTTTGTTTTAGAGCTAGAAATAGCAAGTTAAAATAAGGCTAGTCCGTTTTTAGCGCTTGCGCCAATTCTGCAGACAAATGGCTAT
CTTTACATGTAGCTTGTGCATTGAATCCAATTATAATTTGCCTTGGCACCAGCTGAGCCAGACAAGAAAGAAAGCTTCCCAGAAGTATA
TCGATTTAGAAGGGTTGACGTCACTTGCTGACTGCACTAATACAGCAAATGATGCAATTAGAATGATTCAAGTGAAATTCCCAAATTAC
TGATTTTTCTCTGGATTTGGTTATCAGATTACATTCGAAGCTAAGATTAGCTACCGAAATTGTCGATCAAATCAGGAAATCCTTTCTCT
ATCGAAAAAGGCATTCGCACATCTTCCTCTCTATGCCATATACACGAAGGGTAGGTACATTGACGTCTTTGCCAGAAGTTGAACTGCAT
CGTTCAAGGTACAGAATGAACGACTAACAGACACAAGCACGTTTTGCTGTCCATTCAGACACAGGGATGGTACCCATATTCGATCGATA
TAGAGCCATCCAACCGAACAGAGGTATATGTATGAATGTATTGCTGAAATTTTCTAGAAGTACAACCACCACTACGACAGTGTCTATAA
AACGCCCCTGCAAAGGCGAAACCAGCTCAATCGAATACGTTTCCTAGTGGAGTGAACATTACGCGGCCCAAGTAAGCAGTGCCAGTGCA
AGTGAAGTGAAGTCTCTAGTGAAAAAGAGTGATCCAATTAGCCAGAGGAGAAAATTTCAGAGTGAACAAAGCTTTATTCAAAGGACAAT
TACTATTAAATTGGTGAAAGTGCATTTCGGTGAAGGGAATCTTCTAGTGAAGGTAGGTAAATTAATTGATGAAATTATAGCTATGAGCG
AAAACTAGTTTGGTGAATGATTCCTTTGTCTTTGAATGAGCAAACTATTTTCCAAGATGGCGACTATTGAGCTTTGAGTGATTAGTGAA
AATTTGCAACGCAGTTTCATCATCATTGATAAAACCCAATTGTGATTCACAGCGATAATCATATTTCGTTGAATCATCGCTACTAATTG
AATTAAATTTCTAGAATAATAAGAATAACGTATTTGCTCCGTCACATATCTAAAATAAATATTTTGATGGTTAATTACCCATTAAGGTA
ATATTAACACATATCGAGAAAAACCTTGAGGAAATCGTGAAAACTTGAAGATACGCAATTTCAAAACTACGTAGTTCAAAGTCGAAAAC
AAGTTAATTTTTCACTTAAAAGTAGGGCGTTGTTGTGACGTCATCACCTTCAAGTGTATATTTTTCACTTGGCCTGCGACTGCAAACGC
AGACAAAGCAAAACAAGTTTAAAACCTGTCGTGTCGTGCTCGAAGCCAAAGGCAATGAATCAATATCAAATGAGAGTTTGCATTTCACA
ACCAATTACTCAAGCGTTTCCTCGTTTCTTTTTCTGCTCAACAGAGATTTCAACACCAAGCTCGATAAGCTTGTTCGAATCTCGAGTGC
GCGCTTCCGGAGGTATACACCTAGGCGGTACCACTGCAGTGAATTCGGAGCTCTACCGGTGCCACCATGGACTATAAGGACCACGACGG
AGACTACAAGGATCATGATATTGATTACAAAGACGATGACGATAAGATGGCCCCAAAGAAGAAGCGGAAGGTCGGTATCCACGGAGTCC
CAGCAGCCGACAAGAAGTACAGCATCGGCCTGGACATCGGCACCAACTCTGTGGGCTGGGCCGTGATCACCGACGAGTACAAGGTGCCC
AGCAAGAAATTCAAGGTGCTGGGCAACACCGACCGGCACAGCATCAAGAAGAACCTGATCGGAGCCCTGCTGTTCGACAGCGGCGAAAC
AGCCGAGGCCACCCGGCTGAAGAGAACCGCCAGAAGAAGATACACCAGACGGAAGAACCGGATCTGCTATCTGCAAGAGATCTTCAGCA
ACGAGATGGCCAAGGTGGACGACAGCTTCTTCCACAGACTGGAAGAGTCCTTCCTGGTGGAAGAGGATAAGAAGCACGAGCGGCACCCC
ATCTTCGGCAACATCGTGGACGAGGTGGCCTACCACGAGAAGTACCCCACCATCTACCACCTGAGAAAGAAACTGGTGGACAGCACCGA
CAAGGCCGACCTGCGGCTGATCTATCTGGCCCTGGCCCACATGATCAAGTTCCGGGGCCACTTCCTGATCGAGGGCGACCTGAACCCCG
ACAACAGCGACGTGGACAAGCTGTTCATCCAGCTGGTGCAGACCTACAACCAGCTGTTCGAGGAAAACCCCATCAACGCCAGCGGCGTG
GACGCCAAGGCCATCCTGTCTGCCAGACTGAGCAAGAGCAGACGGCTGGAAAATCTGATCGCCCAGCTGCCCGGCGAGAAGAAGAATGG
CCTGTTCGGAAACCTGATTGCCCTGAGCCTGGGCCTGACCCCCAACTTCAAGAGCAACTTCGACCTGGCCGAGGATGCCAAACTGCAGC
TGAGCAAGGACACCTACGACGACGACCTGGACAACCTGCTGGCCCAGATCGGCGACCAGTACGCCGACCTGTTTCTGGCCGCCAAGAAC
CTGTCCGACGCCATCCTGCTGAGCGACATCCTGAGAGTGAACACCGAGATCACCAAGGCCCCCCTGAGCGCCTCTATGATCAAGAGATA
CGACGAGCACCACCAGGACCTGACCCTGCTGAAAGCTCTCGTGCGGCAGCAGCTGCCTGAGAAGTACAAAGAGATTTTCTTCGACCAGA
GCAAGAACGGCTACGCCGGCTACATTGACGGCGGAGCCAGCCAGGAAGAGTTCTACAAGTTCATCAAGCCCATCCTGGAAAAGATGGAC
GGCACCGAGGAACTGCTCGTGAAGCTGAACAGAGAGGACCTGCTGCGGAAGCAGCGGACCTTCGACAACGGCAGCATCCCCCACCAGAT
CCACCTGGGAGAGCTGCACGCCATTCTGCGGCGGCAGGAAGATTTTTACCCATTCCTGAAGGACAACCGGGAAAAGATCGAGAAGATCC
TGACCTTCCGCATCCCCTACTACGTGGGCCCTCTGGCCAGGGGAAACAGCAGATTCGCCTGGATGACCAGAAAGAGCGAGGAAACCATC
ACCCCCTGGAACTTCGAGGAAGTGGTGGACAAGGGCGCTTCCGCCCAGAGCTTCATCGAGCGGATGACCAACTTCGATAAGAACCTGCC
CAACGAGAAGGTGCTGCCCAAGCACAGCCTGCTGTACGAGTACTTCACCGTGTATAACGAGCTGACCAAAGTGAAATACGTGACCGAGG
GAATGAGAAAGCCCGCCTTCCTGAGCGGCGAGCAGAAAAAGGCCATCGTGGACCTGCTGTTCAAGACCAACCGGAAAGTGACCGTGAAG
CAGCTGAAAGAGGACTACTTCAAGAAAATCGAGTGCTTCGACTCCGTGGAAATCTCCGGCGTGGAAGATCGGTTCAACGCCTCCCTGGG
CACATACCACGATCTGCTGAAAATTATCAAGGACAAGGACTTCCTGGACAATGAGGAAAACGAGGACATTCTGGAAGATATCGTGCTGA
CCCTGACACTGTTTGAGGACAGAGAGATGATCGAGGAACGGCTGAAAACCTATGCCCACCTGTTCGACGACAAAGTGATGAAGCAGCTG
AAGCGGCGGAGATACACCGGCTGGGGCAGGCTGAGCCGGAAGCTGATCAACGGCATCCGGGACAAGCAGTCCGGCAAGACAATCCTGGA
TTTCCTGAAGTCCGACGGCTTCGCCAACAGAAACTTCATGCAGCTGATCCACGACGACAGCCTGACCTTTAAAGAGGACATCCAGAAAG
CCCAGGTGTCCGGCCAGGGCGATAGCCTGCACGAGCACATTGCCAATCTGGCCGGCAGCCCCGCCATTAAGAAGGGCATCCTGCAGACA
GTGAAGGTGGTGGACGAGCTCGTGAAAGTGATGGGCCGGCACAAGCCCGAGAACATCGTGATCGAAATGGCCAGAGAGAACCAGACCAC
CCAGAAGGGACAGAAGAACAGCCGCGAGAGAATGAAGCGGATCGAAGAGGGCATCAAAGAGCTGGGCAGCCAGATCCTGAAAGAACACC
CCGTGGAAAACACCCAGCTGCAGAACGAGAAGCTGTACCTGTACTACCTGCAGAATGGGCGGGATATGTACGTGGACCAGGAACTGGAC
ATCAACCGGCTGTCCGACTACGATGTGGACCATATCGTGCCTCAGAGCTTTCTGAAGGACGACTCCATCGACAACAAGGTGCTGACCAG
AAGCGACAAGAACCGGGGCAAGAGCGACAACGTGCCCTCCGAAGAGGTCGTGAAGAAGATGAAGAACTACTGGCGGCAGCTGCTGAACG
CCAAGCTGATTACCCAGAGAAAGTTCGACAATCTGACCAAGGCCGAGAGAGGCGGCCTGAGCGAACTGGATAAGGCCGGCTTCATCAAG
AGACAGCTGGTGGAAACCCGGCAGATCACAAAGCACGTGGCACAGATCCTGGACTCCCGGATGAACACTAAGTACGACGAGAATGACAA
GCTGATCCGGGAAGTGAAAGTGATCACCCTGAAGTCCAAGCTGGTGTCCGATTTCCGGAAGGATTTCCAGTTTTACAAAGTGCGCGAGA

TCAACAACTACCACCACGCCCACGACGCCTACCTGAACGCCGTCGTGGGAACCGCCCTGATCAAAAAGTACCCTAAGCTGGAAAGCGAG
TTCGTGTACGGCGACTACAAGGTGTACGACGTGCGGAAGATGATCGCCAAGAGCGAGCAGGAAATCGGCAAGGCTACCGCCAAGTACTT
CTTCTACAGCAACATCATGAACTTTTTCAAGACCGAGATTACCCTGGCCAACGGCGAGATCCGGAAGCGGCCTCTGATCGAGACAAACG
GCGAAACCGGGGAGATCGTGTGGGATAAGGGCCGGGATTTTGCCACCGTGCGGAAAGTGCTGAGCATGCCCCAAGTGAATATCGTGAAA
AAGACCGAGGTGCAGACAGGCGGCTTCAGCAAAGAGTCTATCCTGCCCAAGAGGAACAGCGATAAGCTGATCGCCAGAAAGAAGGACTG
GGACCCTAAGAAGTACGGCGGCTTCGACAGCCCCACCGTGGCCTATTCTGTGCTGGTGGTGGCCAAAGTGGAAAAGGGCAAGTCCAAGA
AACTGAAGAGTGTGAAAGAGCTGCTGGGGATCACCATCATGGAAAGAAGCAGCTTCGAGAAGAATCCCATCGACTTTCTGGAAGCCAAG
GGCTACAAAGAAGTGAAAAAGGACCTGATCATCAAGCTGCCTAAGTACTCCCTGTTCGAGCTGGAAAACGGCCGGAAGAGAATGCTGGC
CTCTGCCGGCGAACTGCAGAAGGGAAACGAACTGGCCCTGCCCTCCAAATATGTGAACTTCCTGTACCTGGCCAGCCACTATGAGAAGC
TGAAGGGCTCCCCCGAGGATAATGAGCAGAAACAGCTGTTTGTGGAACAGCACAAGCACTACCTGGACGAGATCATCGAGCAGATCAGC
GAGTTCTCCAAGAGAGTGATCCTGGCCGACGCTAATCTGGACAAAGTGCTGTCCGCCTACAACAAGCACCGGGATAAGCCCATCAGAGA
GCAGGCCaAGAATATCATCCACCTGTTTACCCTGACCAATCTGGGAGCCCCTGCCGCCTTCAAGTACTTTGACACCACCATCGACCGGA
AGAGGTACACCAGCACCAAAGAGGTGCTGGACGCCACCCTGATCCACCAGAGCATCACCGGCCTGTACGAGACACGGATCGACCTGTCT
CAGCTGGGAGGCGACAGCCCCAAGAAGAAGAGAAAGGTGGAGGCCAGCTAATAGGACCCAGCTTTCTTGTACAAAGTGGTGACGTAAGC
TAGCAGGATCTTTGTGAAGGAACCTTACTTCTGTGGTGTGACATAATTGGACAAACTACCTACAGAGATTTAAAGCTCTAAGGTAAATA
TAAAATTTTTAAGTGTATAATGTGTTAAACTACTGATTCTAATTGTTTGTGTATTTTAGATTCCAACCTATGGAACTGATGAATGGGAG
CAGTGGTGGAATGCCTTTAATGAGGAAAACCTGTTTTGCTCAGAAGAAATGCCATCTAGTGATGATGAGGCTACTGCTGACTCTCAACA
TTCTACTCCTCCAAAAAAGAAGAGAAAGGTAGttGACCCCAAGGACTTTCCTTCAGAATTGCTAAGTTTTTTGAGTCATGCTGTGTTTA
GTAATAGAACTCTTGCTTGCTTTGCTATTTACACCACAAAGGAAAAAGCTGCACTGCTATACAAGAAAATTATGGAAAAATATTCTGTA
ACCTTTATAAGTAGGCATAACAGTTATAATCATAACATACTGTTTTTTCTTACTCCACACAGGCATAGAGTGTCTGCTATTAATAACTA
TGCTCAAAAATTGTGTACCTTTAGCTTTTTAATTTGTAAAGGGGTTAATAAGGAATATTTGATGTATAGTGCCTTGACTAGAGATCATA
ATCAGCCATACCACATTTGTAGAGGTTTTACTTGCTTTAAAAAACCTCCCACACCTCCCCCTGAACCTGAAACATAAAATGAATGCAAT
TGTTGTTGTTAACTTGTTTATTGCAGCTTATAATGGTTACAAATAAAGCAATAGCATCACAAATTTCACAAATAAAGCATTTTTTTCAC
TGCATTCTAGTTGTGGTTTGTCCAAACTCATCAATGTATCTTATCATGTCTGGATCCCGTTTAAACTACGCGTAATTCAAACAGGGTTC
TGGCGTCGTTCTCGTACTGTTTTCCCCAGGCCAGTGCTTTAGCGTTATTGAAAAAGGAAGAGTATGAGTATTCAACATTTCCGTGTCGC
CCTTATTCCCTTTTTTGCGGCATTTTGCCTTCCTGTTTTTGCTCACCCAGAAACGCTGGTGAAAGTAAAAGATGCTGAAGATCAGTTGG
GTGCACGAGTGGGTTACATCGAACTGGATCTCAACAGCGGTAAGATCCTTGAGAGTTTTCGCCCCGAAGAACGTTTTCCAATGATGAGC
ACTTTTAAAGTTCTGCTATGTGGCGCGGTATTATCCCGTATTGACGCCGGGCAAGAGCAACTCGGTCGCCGCATACACTATTCTCAGAA
TGACTTGGTTGAGTACTCACCAGTCACAGAAAAGCATCTTACGGATGGCATGACAGTAAGAGAATTATGCAGTGCTGCCATAACCATGA
GTGATAACACTGCGGCCAACTTACTTCTGACAACGATCGGAGGACCGAAGGAGCTAACCGCTTTTTTGCACAACATGGGGGATCATGTA
ACTCGCCTTGATCGTTGGGAACCGGAGCTGAATGAAGCCATACCAAACGACGAGCGTGACACCACGATGCCTGTAGCAATGGCAACAAC
GTTGCGCAAACTATTAACTGGCGAACTACTTACTCTAGCTTCCCGGCAACAATTAATAGACTGGATGGAGGCGGATAAAGTTGCAGGAC
CACTTCTGCGCTCGGCCCTTCCGGCTGGCTGGTTTATTGCTGATAAATCTGGAGCCGGTGAGCGTGGGTCTCGCGGTATCATTGCAGCA
CTGGGGCCAGATGGTAAGCCCTCCCGTATCGTAGTTATCTACACGACGGGGAGTCAGGCAACTATGGATGAACGAAATAGACAGATCGC
TGAGATAGGTGCCTCACTGATTAAGCATTGGTAACTGTCAGACCAAGTTTACTCATATATACTTTAGATTGATTTAAAACTTCATTTTT
AATTTAAAAGGATCTAGGTGAAGATCCTTTTTGATAATCTCATGACCAAAATCCCTTAACGTGAGTTTTCGTTCCACTGAGCGTCAGAC
CCCGTAGAAAAGATCAAAGGATCTTCTTGAGATCCTTTTTTTCTGCGCGTAATCTGCTGCTTGCAAACAAAAAAACCACCGCTACCAGC
GGTGGTTTGTTTGCCGGATCAAGAGCTACCAACTCTTTTTCCGAAGGTAACTGGCTTCAGCAGAGCGCAGATACCAAATACTGTTCTTC
TAGTGTAGCCGTAGTTAGGCCACCACTTCAAGAACTCTGTAGCACCGCCTACATACCTCGCTCTGCTAATCCTGTTACCAGTGGCTGCT
GCCAGTGGCGATAAGTCGTGTCTTACCGGGTTGGACTCAAGACGATAGTTACCGGATAAGGCGCAGCGGTCGGGCTGAACGGGGGGTTC
GTGCACACAGCCCAGCTTGGAGCGAACGACCTACACCGAACTGAGATACCTACAGCGTGAGCTATGAGAAAGCGCCACGCTTCCCGAAG
GGAGAAAGGCGGACAGGTATCCGGTAAGCGGCAGGGTCGGAACAGGAGAGCGCACGAGGGAGCTTCCAGGGGGAAACGCCTGGTATCTT
TATAGTCCTGTCGGGTTTCGCCACCTCTGACTTGAGCGTCGATTTTTGTGATGCTCGTCAGGGGGGCGGAGCCTATGGAAAAACGCCAG
CAACGCGGCCTTTTTACGGTTCCTGGCCTTTTGCTGGCCTTTTGCTCACATGTTCTTTCCTGCGTTATCCCCTGATTCTGTGGATAACC
GTATTACCGCCTTTGAGTGAGCTGATACCGCTCGCCGCAGCCGAACGACCGAGCGCAGCGAGTCAGTGAGCGAGGAAGCGGAAGA

pKRG3-mU6-PUb-hSpCas9

*aae* U6
*BbsI* sites for sgRNA cloning
sgRNA tracrRNA and U6 terminator
*aae* PUb promoter
NLS
hSpCas9

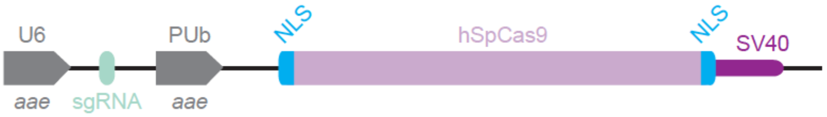


GCGCCCAATACGCAAACCGCCTCTCCCCGCGGGTTGGCCGATTCATTAATGCAGGCAACTCGTGAAAGGTAGGCGGATCAGCGAATGAA
ATCGCCCATCGAGTTGATACGTCCATCCATCGCTAGAACCGCGTTCGCTGTAGAAGACTATATAAGAGCAGAGGCAAGAGTAGTGAAAT
gGAGACGaCGTCTCtGTTTTAGAGCTAGAAATAGCAAGTTAAAATAAGGCTAGTCCGTTATCAACTTGAAAAAGTGGCACCGAGTCGGT
GCTTTTTTGTTTTAGAGCTAGAAATAGCAAGTTAAAATAAGGCTAGTCCGTTTTTAGCGCTTGCGCCAATTCTGCAGACAAATGGCTAT
CTTTACATGTAGCTTGTGCATTGAATCCAATTATAATTTGCCTTGGCACCAGCTGAGCCAGACAAGAAAGAAAGCTTCCCAGAAGTATA
TCGATTTAGAAGGGTTGACGTCACTTGCTGACTGCACTAATACAGCAAATGATGCAATTAGAATGATTCAAGTGAAATTCCCAAATTAC
TGATTTTTCTCTGGATTTGGTTATCAGATTACATTCGAAGCTAAGATTAGCTACCGAAATTGTCGATCAAATCAGGAAATCCTTTCTCT
ATCGAAAAAGGCATTCGCACATCTTCCTCTCTATGCCATATACACGAAGGGTAGGTACATTGACGTCTTTGCCAGAAGTTGAACTGCAT
CGTTCAAGGTACAGAATGAACGACTAACAGACACAAGCACGTTTTGCTGTCCATTCAGACACAGGGATGGTACCCATATTCGATCGATA
TAGAGCCATCCAACCGAACAGAGGTATATGTATGAATGTATTGCTGAAATTTTCTAGAAGTACAACCACCACTACGACAGTGTCTATAA
AACGCCCCTGCAAAGGCGAAACCAGCTCAATCGAATACGTTTCCTAGTGGAGTGAACATTACGCGGCCCAAGTAAGCAGTGCCAGTGCA
AGTGAAGTGAAGTCTCTAGTGAAAAAGAGTGATCCAATTAGCCAGAGGAGAAAATTTCAGAGTGAACAAAGCTTTATTCAAAGGACAAT
TACTATTAAATTGGTGAAAGTGCATTTCGGTGAAGGGAATCTTCTAGTGAAGGTAGGTAAATTAATTGATGAAATTATAGCTATGAGCG
AAAACTAGTTTGGTGAATGATTCCTTTGTCTTTGAATGAGCAAACTATTTTCCAAGATGGCGACTATTGAGCTTTGAGTGATTAGTGAA
AATTTGCAACGCAGTTTCATCATCATTGATAAAACCCAATTGTGATTCACAGCGATAATCATATTTCGTTGAATCATCGCTACTAATTG
AATTAAATTTCTAGAATAATAAGAATAACGTATTTGCTCCGTCACATATCTAAAATAAATATTTTGATGGTTAATTACCCATTAAGGTA
ATATTAACACATATCGAGAAAAACCTTGAGGAAATCGTGAAAACTTGAAGATACGCAATTTCAAAACTACGTAGTTCAAAGTCGAAAAC
AAGTTAATTTTTCACTTAAAAGTAGGGCGTTGTTGTGACGTCATCACCTTCAAGTGTATATTTTTCACTTGGCCTGCGACTGCAAACGC
AGACAAAGCAAAACAAGTTTAAAACCTGTCGTGTCGTGCTCGAAGCCAAAGGCAATGAATCAATATCAAATGAGAGTTTGCATTTCACA
ACCAATTACTCAAGCGTTTCCTCGTTTCTTTTTCTGCTCAACAGAGATTTCAACACCAAGCTCGATAAGCTTGTTCGAATCTCGAGTGC
GCGCTTCCGGAGGTATACACCTAGGCGGTACCACTGCAGTGAATTCGGAGCTCTACCGGTGCCACCATGGCCCCAAAGAAGAAGCGGAA
GGTCGGTATCCACGGAGTCCCAGCAGCCGACAAGAAGTACAGCATCGGCCTGGACATCGGCACCAACTCTGTGGGCTGGGCCGTGATCA
CCGACGAGTACAAGGTGCCCAGCAAGAAATTCAAGGTGCTGGGCAACACCGACCGGCACAGCATCAAGAAGAACCTGATCGGAGCCCTG
CTGTTCGACAGCGGCGAAACAGCCGAGGCCACCCGGCTGAAGAGAACCGCCAGAAGAAGATACACCAGACGGAAGAACCGGATCTGCTA
TCTGCAAGAGATCTTCAGCAACGAGATGGCCAAGGTGGACGACAGCTTCTTCCACAGACTGGAAGAGTCCTTCCTGGTGGAAGAGGATA
AGAAGCACGAGCGGCACCCCATCTTCGGCAACATCGTGGACGAGGTGGCCTACCACGAGAAGTACCCCACCATCTACCACCTGAGAAAG
AAACTGGTGGACAGCACCGACAAGGCCGACCTGCGGCTGATCTATCTGGCCCTGGCCCACATGATCAAGTTCCGGGGCCACTTCCTGAT
CGAGGGCGACCTGAACCCCGACAACAGCGACGTGGACAAGCTGTTCATCCAGCTGGTGCAGACCTACAACCAGCTGTTCGAGGAAAACC
CCATCAACGCCAGCGGCGTGGACGCCAAGGCCATCCTGTCTGCCAGACTGAGCAAGAGCAGACGGCTGGAAAATCTGATCGCCCAGCTG
CCCGGCGAGAAGAAGAATGGCCTGTTCGGAAACCTGATTGCCCTGAGCCTGGGCCTGACCCCCAACTTCAAGAGCAACTTCGACCTGGC
CGAGGATGCCAAACTGCAGCTGAGCAAGGACACCTACGACGACGACCTGGACAACCTGCTGGCCCAGATCGGCGACCAGTACGCCGACC
TGTTTCTGGCCGCCAAGAACCTGTCCGACGCCATCCTGCTGAGCGACATCCTGAGAGTGAACACCGAGATCACCAAGGCCCCCCTGAGC
GCCTCTATGATCAAGAGATACGACGAGCACCACCAGGACCTGACCCTGCTGAAAGCTCTCGTGCGGCAGCAGCTGCCTGAGAAGTACAA
AGAGATTTTCTTCGACCAGAGCAAGAACGGCTACGCCGGCTACATTGACGGCGGAGCCAGCCAGGAAGAGTTCTACAAGTTCATCAAGC
CCATCCTGGAAAAGATGGACGGCACCGAGGAACTGCTCGTGAAGCTGAACAGAGAGGACCTGCTGCGGAAGCAGCGGACCTTCGACAAC
GGCAGCATCCCCCACCAGATCCACCTGGGAGAGCTGCACGCCATTCTGCGGCGGCAGGAAGATTTTTACCCATTCCTGAAGGACAACCG
GGAAAAGATCGAGAAGATCCTGACCTTCCGCATCCCCTACTACGTGGGCCCTCTGGCCAGGGGAAACAGCAGATTCGCCTGGATGACCA
GAAAGAGCGAGGAAACCATCACCCCCTGGAACTTCGAGGAAGTGGTGGACAAGGGCGCTTCCGCCCAGAGCTTCATCGAGCGGATGACC
AACTTCGATAAGAACCTGCCCAACGAGAAGGTGCTGCCCAAGCACAGCCTGCTGTACGAGTACTTCACCGTGTATAACGAGCTGACCAA
AGTGAAATACGTGACCGAGGGAATGAGAAAGCCCGCCTTCCTGAGCGGCGAGCAGAAAAAGGCCATCGTGGACCTGCTGTTCAAGACCA
ACCGGAAAGTGACCGTGAAGCAGCTGAAAGAGGACTACTTCAAGAAAATCGAGTGCTTCGACTCCGTGGAAATCTCCGGCGTGGAAGAT
CGGTTCAACGCCTCCCTGGGCACATACCACGATCTGCTGAAAATTATCAAGGACAAGGACTTCCTGGACAATGAGGAAAACGAGGACAT
TCTGGAAGATATCGTGCTGACCCTGACACTGTTTGAGGACAGAGAGATGATCGAGGAACGGCTGAAAACCTATGCCCACCTGTTCGACG
ACAAAGTGATGAAGCAGCTGAAGCGGCGGAGATACACCGGCTGGGGCAGGCTGAGCCGGAAGCTGATCAACGGCATCCGGGACAAGCAG
TCCGGCAAGACAATCCTGGATTTCCTGAAGTCCGACGGCTTCGCCAACAGAAACTTCATGCAGCTGATCCACGACGACAGCCTGACCTT
TAAAGAGGACATCCAGAAAGCCCAGGTGTCCGGCCAGGGCGATAGCCTGCACGAGCACATTGCCAATCTGGCCGGCAGCCCCGCCATTA
AGAAGGGCATCCTGCAGACAGTGAAGGTGGTGGACGAGCTCGTGAAAGTGATGGGCCGGCACAAGCCCGAGAACATCGTGATCGAAATG
GCCAGAGAGAACCAGACCACCCAGAAGGGACAGAAGAACAGCCGCGAGAGAATGAAGCGGATCGAAGAGGGCATCAAAGAGCTGGGCAG
CCAGATCCTGAAAGAACACCCCGTGGAAAACACCCAGCTGCAGAACGAGAAGCTGTACCTGTACTACCTGCAGAATGGGCGGGATATGT
ACGTGGACCAGGAACTGGACATCAACCGGCTGTCCGACTACGATGTGGACCATATCGTGCCTCAGAGCTTTCTGAAGGACGACTCCATC
GACAACAAGGTGCTGACCAGAAGCGACAAGAACCGGGGCAAGAGCGACAACGTGCCCTCCGAAGAGGTCGTGAAGAAGATGAAGAACTA
CTGGCGGCAGCTGCTGAACGCCAAGCTGATTACCCAGAGAAAGTTCGACAATCTGACCAAGGCCGAGAGAGGCGGCCTGAGCGAACTGG
ATAAGGCCGGCTTCATCAAGAGACAGCTGGTGGAAACCCGGCAGATCACAAAGCACGTGGCACAGATCCTGGACTCCCGGATGAACACT
AAGTACGACGAGAATGACAAGCTGATCCGGGAAGTGAAAGTGATCACCCTGAAGTCCAAGCTGGTGTCCGATTTCCGGAAGGATTTCCA
GTTTTACAAAGTGCGCGAGATCAACAACTACCACCACGCCCACGACGCCTACCTGAACGCCGTCGTGGGAACCGCCCTGATCAAAAAGT
ACCCTAAGCTGGAAAGCGAGTTCGTGTACGGCGACTACAAGGTGTACGACGTGCGGAAGATGATCGCCAAGAGCGAGCAGGAAATCGGC

AAGGCTACCGCCAAGTACTTCTTCTACAGCAACATCATGAACTTTTTCAAGACCGAGATTACCCTGGCCAACGGCGAGATCCGGAAGCG
GCCTCTGATCGAGACAAACGGCGAAACCGGGGAGATCGTGTGGGATAAGGGCCGGGATTTTGCCACCGTGCGGAAAGTGCTGAGCATGC
CCCAAGTGAATATCGTGAAAAAGACCGAGGTGCAGACAGGCGGCTTCAGCAAAGAGTCTATCCTGCCCAAGAGGAACAGCGATAAGCTG
ATCGCCAGAAAGAAGGACTGGGACCCTAAGAAGTACGGCGGCTTCGACAGCCCCACCGTGGCCTATTCTGTGCTGGTGGTGGCCAAAGT
GGAAAAGGGCAAGTCCAAGAAACTGAAGAGTGTGAAAGAGCTGCTGGGGATCACCATCATGGAAAGAAGCAGCTTCGAGAAGAATCCCA
TCGACTTTCTGGAAGCCAAGGGCTACAAAGAAGTGAAAAAGGACCTGATCATCAAGCTGCCTAAGTACTCCCTGTTCGAGCTGGAAAAC
GGCCGGAAGAGAATGCTGGCCTCTGCCGGCGAACTGCAGAAGGGAAACGAACTGGCCCTGCCCTCCAAATATGTGAACTTCCTGTACCT
GGCCAGCCACTATGAGAAGCTGAAGGGCTCCCCCGAGGATAATGAGCAGAAACAGCTGTTTGTGGAACAGCACAAGCACTACCTGGACG
AGATCATCGAGCAGATCAGCGAGTTCTCCAAGAGAGTGATCCTGGCCGACGCTAATCTGGACAAAGTGCTGTCCGCCTACAACAAGCAC
CGGGATAAGCCCATCAGAGAGCAGGCCaAGAATATCATCCACCTGTTTACCCTGACCAATCTGGGAGCCCCTGCCGCCTTCAAGTACTT
TGACACCACCATCGACCGGAAGAGGTACACCAGCACCAAAGAGGTGCTGGACGCCACCCTGATCCACCAGAGCATCACCGGCCTGTACG
AGACACGGATCGACCTGTCTCAGCTGGGAGGCGACAGCCCCAAGAAGAAGAGAAAGGTGGAGGCCAGCTAATAGGACCCAGCTTTCTTG
TACAAAGTGGTGACGTAAGCTAGCAGGATCTTTGTGAAGGAACCTTACTTCTGTGGTGTGACATAATTGGACAAACTACCTACAGAGAT
TTAAAGCTCTAAGGTAAATATAAAATTTTTAAGTGTATAATGTGTTAAACTACTGATTCTAATTGTTTGTGTATTTTAGATTCCAACCT
ATGGAACTGATGAATGGGAGCAGTGGTGGAATGCCTTTAATGAGGAAAACCTGTTTTGCTCAGAAGAAATGCCATCTAGTGATGATGAG
GCTACTGCTGACTCTCAACATTCTACTCCTCCAAAAAAGAAGAGAAAGGTAGttGACCCCAAGGACTTTCCTTCAGAATTGCTAAGTTT
TTTGAGTCATGCTGTGTTTAGTAATAGAACTCTTGCTTGCTTTGCTATTTACACCACAAAGGAAAAAGCTGCACTGCTATACAAGAAAA
TTATGGAAAAATATTCTGTAACCTTTATAAGTAGGCATAACAGTTATAATCATAACATACTGTTTTTTCTTACTCCACACAGGCATAGA
GTGTCTGCTATTAATAACTATGCTCAAAAATTGTGTACCTTTAGCTTTTTAATTTGTAAAGGGGTTAATAAGGAATATTTGATGTATAG
TGCCTTGACTAGAGATCATAATCAGCCATACCACATTTGTAGAGGTTTTACTTGCTTTAAAAAACCTCCCACACCTCCCCCTGAACCTG
AAACATAAAATGAATGCAATTGTTGTTGTTAACTTGTTTATTGCAGCTTATAATGGTTACAAATAAAGCAATAGCATCACAAATTTCAC
AAATAAAGCATTTTTTTCACTGCATTCTAGTTGTGGTTTGTCCAAACTCATCAATGTATCTTATCATGTCTGGATCCCGTTTAAACTAC
GCGTAATTCAAACAGGGTTCTGGCGTCGTTCTCGTACTGTTTTCCCCAGGCCAGTGCTTTAGCGTTATTGAAAAAGGAAGAGTATGAGT
ATTCAACATTTCCGTGTCGCCCTTATTCCCTTTTTTGCGGCATTTTGCCTTCCTGTTTTTGCTCACCCAGAAACGCTGGTGAAAGTAAA
AGATGCTGAAGATCAGTTGGGTGCACGAGTGGGTTACATCGAACTGGATCTCAACAGCGGTAAGATCCTTGAGAGTTTTCGCCCCGAAG
AACGTTTTCCAATGATGAGCACTTTTAAAGTTCTGCTATGTGGCGCGGTATTATCCCGTATTGACGCCGGGCAAGAGCAACTCGGTCGC
CGCATACACTATTCTCAGAATGACTTGGTTGAGTACTCACCAGTCACAGAAAAGCATCTTACGGATGGCATGACAGTAAGAGAATTATG
CAGTGCTGCCATAACCATGAGTGATAACACTGCGGCCAACTTACTTCTGACAACGATCGGAGGACCGAAGGAGCTAACCGCTTTTTTGC
ACAACATGGGGGATCATGTAACTCGCCTTGATCGTTGGGAACCGGAGCTGAATGAAGCCATACCAAACGACGAGCGTGACACCACGATG
CCTGTAGCAATGGCAACAACGTTGCGCAAACTATTAACTGGCGAACTACTTACTCTAGCTTCCCGGCAACAATTAATAGACTGGATGGA
GGCGGATAAAGTTGCAGGACCACTTCTGCGCTCGGCCCTTCCGGCTGGCTGGTTTATTGCTGATAAATCTGGAGCCGGTGAGCGTGGGT
CTCGCGGTATCATTGCAGCACTGGGGCCAGATGGTAAGCCCTCCCGTATCGTAGTTATCTACACGACGGGGAGTCAGGCAACTATGGAT
GAACGAAATAGACAGATCGCTGAGATAGGTGCCTCACTGATTAAGCATTGGTAACTGTCAGACCAAGTTTACTCATATATACTTTAGAT
TGATTTAAAACTTCATTTTTAATTTAAAAGGATCTAGGTGAAGATCCTTTTTGATAATCTCATGACCAAAATCCCTTAACGTGAGTTTT
CGTTCCACTGAGCGTCAGACCCCGTAGAAAAGATCAAAGGATCTTCTTGAGATCCTTTTTTTCTGCGCGTAATCTGCTGCTTGCAAACA
AAAAAACCACCGCTACCAGCGGTGGTTTGTTTGCCGGATCAAGAGCTACCAACTCTTTTTCCGAAGGTAACTGGCTTCAGCAGAGCGCA
GATACCAAATACTGTTCTTCTAGTGTAGCCGTAGTTAGGCCACCACTTCAAGAACTCTGTAGCACCGCCTACATACCTCGCTCTGCTAA
TCCTGTTACCAGTGGCTGCTGCCAGTGGCGATAAGTCGTGTCTTACCGGGTTGGACTCAAGACGATAGTTACCGGATAAGGCGCAGCGG
TCGGGCTGAACGGGGGGTTCGTGCACACAGCCCAGCTTGGAGCGAACGACCTACACCGAACTGAGATACCTACAGCGTGAGCTATGAGA
AAGCGCCACGCTTCCCGAAGGGAGAAAGGCGGACAGGTATCCGGTAAGCGGCAGGGTCGGAACAGGAGAGCGCACGAGGGAGCTTCCAG
GGGGAAACGCCTGGTATCTTTATAGTCCTGTCGGGTTTCGCCACCTCTGACTTGAGCGTCGATTTTTGTGATGCTCGTCAGGGGGGCGG
AGCCTATGGAAAAACGCCAGCAACGCGGCCTTTTTACGGTTCCTGGCCTTTTGCTGGCCTTTTGCTCACATGTTCTTTCCTGCGTTATC
CCCTGATTCTGTGGATAACCGTATTACCGCCTTTGAGTGAGCTGATACCGCTCGCCGCAGCCGAACGACCGAGCGCAGCGAGTCAGTGA
GCGAGGAAGCGGAAGA

### pKRG3-mU6-PUb-hSpCas9-pAc

*aae* U6
*BbsI* sites for sgRNA cloning
sgRNA tracrRNA and U6 terminator
*aae* PUb promoter
NLS
hSpCas9
T2A
pAc

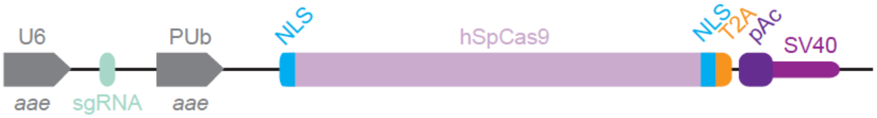


GCGCCCAATACGCAAACCGCCTCTCCCCGCGGGTTGGCCGATTCATTAATGCAGGCAACTCGTGAAAGGTAGGCGGATCAGCGAATGAA
ATCGCCCATCGAGTTGATACGTCCATCCATCGCTAGAACCGCGTTCGCTGTAGAAGACTATATAAGAGCAGAGGCAAGAGTAGTGAAAT
gGAGACGaCGTCTCtGTTTTAGAGCTAGAAATAGCAAGTTAAAATAAGGCTAGTCCGTTATCAACTTGAAAAAGTGGCACCGAGTCGGT
GCTTTTTTGTTTTAGAGCTAGAAATAGCAAGTTAAAATAAGGCTAGTCCGTTTTTAGCGCTTGCGCCAATTCTGCAGACAAATGGCTAT
CTTTACATGTAGCTTGTGCATTGAATCCAATTATAATTTGCCTTGGCACCAGCTGAGCCAGACAAGAAAGAAAGCTTCCCAGAAGTATA
TCGATTTAGAAGGGTTGACGTCACTTGCTGACTGCACTAATACAGCAAATGATGCAATTAGAATGATTCAAGTGAAATTCCCAAATTAC
TGATTTTTCTCTGGATTTGGTTATCAGATTACATTCGAAGCTAAGATTAGCTACCGAAATTGTCGATCAAATCAGGAAATCCTTTCTCT
ATCGAAAAAGGCATTCGCACATCTTCCTCTCTATGCCATATACACGAAGGGTAGGTACATTGACGTCTTTGCCAGAAGTTGAACTGCAT
CGTTCAAGGTACAGAATGAACGACTAACAGACACAAGCACGTTTTGCTGTCCATTCAGACACAGGGATGGTACCCATATTCGATCGATA
TAGAGCCATCCAACCGAACAGAGGTATATGTATGAATGTATTGCTGAAATTTTCTAGAAGTACAACCACCACTACGACAGTGTCTATAA
AACGCCCCTGCAAAGGCGAAACCAGCTCAATCGAATACGTTTCCTAGTGGAGTGAACATTACGCGGCCCAAGTAAGCAGTGCCAGTGCA
AGTGAAGTGAAGTCTCTAGTGAAAAAGAGTGATCCAATTAGCCAGAGGAGAAAATTTCAGAGTGAACAAAGCTTTATTCAAAGGACAAT
TACTATTAAATTGGTGAAAGTGCATTTCGGTGAAGGGAATCTTCTAGTGAAGGTAGGTAAATTAATTGATGAAATTATAGCTATGAGCG
AAAACTAGTTTGGTGAATGATTCCTTTGTCTTTGAATGAGCAAACTATTTTCCAAGATGGCGACTATTGAGCTTTGAGTGATTAGTGAA
AATTTGCAACGCAGTTTCATCATCATTGATAAAACCCAATTGTGATTCACAGCGATAATCATATTTCGTTGAATCATCGCTACTAATTG
AATTAAATTTCTAGAATAATAAGAATAACGTATTTGCTCCGTCACATATCTAAAATAAATATTTTGATGGTTAATTACCCATTAAGGTA
ATATTAACACATATCGAGAAAAACCTTGAGGAAATCGTGAAAACTTGAAGATACGCAATTTCAAAACTACGTAGTTCAAAGTCGAAAAC
AAGTTAATTTTTCACTTAAAAGTAGGGCGTTGTTGTGACGTCATCACCTTCAAGTGTATATTTTTCACTTGGCCTGCGACTGCAAACGC
AGACAAAGCAAAACAAGTTTAAAACCTGTCGTGTCGTGCTCGAAGCCAAAGGCAATGAATCAATATCAAATGAGAGTTTGCATTTCACA
ACCAATTACTCAAGCGTTTCCTCGTTTCTTTTTCTGCTCAACAGAGATTTCAACACCAAGCTCGATAAGCTTGTTCGAATCTCGAGTGC
GCGCTTCCGGAGGTATACACCTAGGCGGTACCACTGCAGTGAATTCGGAGCTCTACCGGTGCCACCATGGCCCCAAAGAAGAAGCGGAA
GGTCGGTATCCACGGAGTCCCAGCAGCCGACAAGAAGTACAGCATCGGCCTGGACATCGGCACCAACTCTGTGGGCTGGGCCGTGATCA
CCGACGAGTACAAGGTGCCCAGCAAGAAATTCAAGGTGCTGGGCAACACCGACCGGCACAGCATCAAGAAGAACCTGATCGGAGCCCTG
CTGTTCGACAGCGGCGAAACAGCCGAGGCCACCCGGCTGAAGAGAACCGCCAGAAGAAGATACACCAGACGGAAGAACCGGATCTGCTA
TCTGCAAGAGATCTTCAGCAACGAGATGGCCAAGGTGGACGACAGCTTCTTCCACAGACTGGAAGAGTCCTTCCTGGTGGAAGAGGATA
AGAAGCACGAGCGGCACCCCATCTTCGGCAACATCGTGGACGAGGTGGCCTACCACGAGAAGTACCCCACCATCTACCACCTGAGAAAG
AAACTGGTGGACAGCACCGACAAGGCCGACCTGCGGCTGATCTATCTGGCCCTGGCCCACATGATCAAGTTCCGGGGCCACTTCCTGAT
CGAGGGCGACCTGAACCCCGACAACAGCGACGTGGACAAGCTGTTCATCCAGCTGGTGCAGACCTACAACCAGCTGTTCGAGGAAAACC
CCATCAACGCCAGCGGCGTGGACGCCAAGGCCATCCTGTCTGCCAGACTGAGCAAGAGCAGACGGCTGGAAAATCTGATCGCCCAGCTG
CCCGGCGAGAAGAAGAATGGCCTGTTCGGAAACCTGATTGCCCTGAGCCTGGGCCTGACCCCCAACTTCAAGAGCAACTTCGACCTGGC
CGAGGATGCCAAACTGCAGCTGAGCAAGGACACCTACGACGACGACCTGGACAACCTGCTGGCCCAGATCGGCGACCAGTACGCCGACC
TGTTTCTGGCCGCCAAGAACCTGTCCGACGCCATCCTGCTGAGCGACATCCTGAGAGTGAACACCGAGATCACCAAGGCCCCCCTGAGC
GCCTCTATGATCAAGAGATACGACGAGCACCACCAGGACCTGACCCTGCTGAAAGCTCTCGTGCGGCAGCAGCTGCCTGAGAAGTACAA
AGAGATTTTCTTCGACCAGAGCAAGAACGGCTACGCCGGCTACATTGACGGCGGAGCCAGCCAGGAAGAGTTCTACAAGTTCATCAAGC
CCATCCTGGAAAAGATGGACGGCACCGAGGAACTGCTCGTGAAGCTGAACAGAGAGGACCTGCTGCGGAAGCAGCGGACCTTCGACAAC
GGCAGCATCCCCCACCAGATCCACCTGGGAGAGCTGCACGCCATTCTGCGGCGGCAGGAAGATTTTTACCCATTCCTGAAGGACAACCG
GGAAAAGATCGAGAAGATCCTGACCTTCCGCATCCCCTACTACGTGGGCCCTCTGGCCAGGGGAAACAGCAGATTCGCCTGGATGACCA
GAAAGAGCGAGGAAACCATCACCCCCTGGAACTTCGAGGAAGTGGTGGACAAGGGCGCTTCCGCCCAGAGCTTCATCGAGCGGATGACC
AACTTCGATAAGAACCTGCCCAACGAGAAGGTGCTGCCCAAGCACAGCCTGCTGTACGAGTACTTCACCGTGTATAACGAGCTGACCAA
AGTGAAATACGTGACCGAGGGAATGAGAAAGCCCGCCTTCCTGAGCGGCGAGCAGAAAAAGGCCATCGTGGACCTGCTGTTCAAGACCA
ACCGGAAAGTGACCGTGAAGCAGCTGAAAGAGGACTACTTCAAGAAAATCGAGTGCTTCGACTCCGTGGAAATCTCCGGCGTGGAAGAT
CGGTTCAACGCCTCCCTGGGCACATACCACGATCTGCTGAAAATTATCAAGGACAAGGACTTCCTGGACAATGAGGAAAACGAGGACAT
TCTGGAAGATATCGTGCTGACCCTGACACTGTTTGAGGACAGAGAGATGATCGAGGAACGGCTGAAAACCTATGCCCACCTGTTCGACG
ACAAAGTGATGAAGCAGCTGAAGCGGCGGAGATACACCGGCTGGGGCAGGCTGAGCCGGAAGCTGATCAACGGCATCCGGGACAAGCAG
TCCGGCAAGACAATCCTGGATTTCCTGAAGTCCGACGGCTTCGCCAACAGAAACTTCATGCAGCTGATCCACGACGACAGCCTGACCTT
TAAAGAGGACATCCAGAAAGCCCAGGTGTCCGGCCAGGGCGATAGCCTGCACGAGCACATTGCCAATCTGGCCGGCAGCCCCGCCATTA
AGAAGGGCATCCTGCAGACAGTGAAGGTGGTGGACGAGCTCGTGAAAGTGATGGGCCGGCACAAGCCCGAGAACATCGTGATCGAAATG
GCCAGAGAGAACCAGACCACCCAGAAGGGACAGAAGAACAGCCGCGAGAGAATGAAGCGGATCGAAGAGGGCATCAAAGAGCTGGGCAG
CCAGATCCTGAAAGAACACCCCGTGGAAAACACCCAGCTGCAGAACGAGAAGCTGTACCTGTACTACCTGCAGAATGGGCGGGATATGT
ACGTGGACCAGGAACTGGACATCAACCGGCTGTCCGACTACGATGTGGACCATATCGTGCCTCAGAGCTTTCTGAAGGACGACTCCATC
GACAACAAGGTGCTGACCAGAAGCGACAAGAACCGGGGCAAGAGCGACAACGTGCCCTCCGAAGAGGTCGTGAAGAAGATGAAGAACTA
CTGGCGGCAGCTGCTGAACGCCAAGCTGATTACCCAGAGAAAGTTCGACAATCTGACCAAGGCCGAGAGAGGCGGCCTGAGCGAACTGG
ATAAGGCCGGCTTCATCAAGAGACAGCTGGTGGAAACCCGGCAGATCACAAAGCACGTGGCACAGATCCTGGACTCCCGGATGAACACT
AAGTACGACGAGAATGACAAGCTGATCCGGGAAGTGAAAGTGATCACCCTGAAGTCCAAGCTGGTGTCCGATTTCCGGAAGGATTTCCA

GTTTTACAAAGTGCGCGAGATCAACAACTACCACCACGCCCACGACGCCTACCTGAACGCCGTCGTGGGAACCGCCCTGATCAAAAAGT
ACCCTAAGCTGGAAAGCGAGTTCGTGTACGGCGACTACAAGGTGTACGACGTGCGGAAGATGATCGCCAAGAGCGAGCAGGAAATCGGC
AAGGCTACCGCCAAGTACTTCTTCTACAGCAACATCATGAACTTTTTCAAGACCGAGATTACCCTGGCCAACGGCGAGATCCGGAAGCG
GCCTCTGATCGAGACAAACGGCGAAACCGGGGAGATCGTGTGGGATAAGGGCCGGGATTTTGCCACCGTGCGGAAAGTGCTGAGCATGC
CCCAAGTGAATATCGTGAAAAAGACCGAGGTGCAGACAGGCGGCTTCAGCAAAGAGTCTATCCTGCCCAAGAGGAACAGCGATAAGCTG
ATCGCCAGAAAGAAGGACTGGGACCCTAAGAAGTACGGCGGCTTCGACAGCCCCACCGTGGCCTATTCTGTGCTGGTGGTGGCCAAAGT
GGAAAAGGGCAAGTCCAAGAAACTGAAGAGTGTGAAAGAGCTGCTGGGGATCACCATCATGGAAAGAAGCAGCTTCGAGAAGAATCCCA
TCGACTTTCTGGAAGCCAAGGGCTACAAAGAAGTGAAAAAGGACCTGATCATCAAGCTGCCTAAGTACTCCCTGTTCGAGCTGGAAAAC
GGCCGGAAGAGAATGCTGGCCTCTGCCGGCGAACTGCAGAAGGGAAACGAACTGGCCCTGCCCTCCAAATATGTGAACTTCCTGTACCT
GGCCAGCCACTATGAGAAGCTGAAGGGCTCCCCCGAGGATAATGAGCAGAAACAGCTGTTTGTGGAACAGCACAAGCACTACCTGGACG
AGATCATCGAGCAGATCAGCGAGTTCTCCAAGAGAGTGATCCTGGCCGACGCTAATCTGGACAAAGTGCTGTCCGCCTACAACAAGCAC
CGGGATAAGCCCATCAGAGAGCAGGCCaAGAATATCATCCACCTGTTTACCCTGACCAATCTGGGAGCCCCTGCCGCCTTCAAGTACTT
TGACACCACCATCGACCGGAAGAGGTACACCAGCACCAAAGAGGTGCTGGACGCCACCCTGATCCACCAGAGCATCACCGGCCTGTACG
AGACACGGATCGACCTGTCTCAGCTGGGAGGCGACAAAAGGCCGGCGGCCACGAAAAAGGCCGGCCAGGCAAAAAAGAAAAAGGGATCA
GGCAAGCTTGAGGGCAGAGGAAGTCTTCTAACATGCGGTGACGTGGAGGAGAATCCCGGCCCTGCTAGCGGTAGCGGCAGCGGTAGCAT
GACCGAGTACAAGCCCACGGTGCGCCTCGCCACCCGCGACGACGTCCCCGGGCCGTACGCACCCTCGCCGCCGCGTTCGCCGACTACC
CCGCCACGCGCCACACCGTCGACCCGGACCGCCACATCGAGCGGGTCACCGAGCTGCAAGAACTCTTCCTCACGCGCGTCGGGCTCGAC
ATCGGCAAGGTGTGGGTCGCGGACGACGGCGCCGCGGTGGCGGTCTGGACCACGCCGGAGAGCGTCGAAGCGGGGGCGGTGTTCGCCGA
GATCGGCCCGCGCATGGCCGAGTTGAGCGGTTCCCGGCTGGCCGCGCAGCAACAGATGGAAGGCCTCCTGGCGCCGCACCGGCCCAAGG
AGCCCGCGTGGTTCCTGGCCACCGTCGGCGTCTCGCCCGACCACCAGGGCAAGGGTCTGGGCAGCGCCGTCGTGCTCCCCGGAGTGGAG
GCGGCCGAGCGCGCCGGGGTGCCCGCCTTCCTGGAGACCTCCGCGCCCCGCAACCTCCCCTTCTACGAGCGGCTCGGCTTCACCGTCAC
CGCCGACGTCGAGGTGCCCGAAGGACCGCGCACCTGGTGCATGACCCGCAAGCCCGGTGCCTAATAGGACCCAGCTTTCTTGTACAAAG
TGGTGACGTAAGCTAGCAGGATCTTTGTGAAGGAACCTTACTTCTGTGGTGTGACATAATTGGACAAACTACCTACAGAGATTTAAAGC
TCTAAGGTAAATATAAAATTTTTAAGTGTATAATGTGTTAAACTACTGATTCTAATTGTTTGTGTATTTTAGATTCCAACCTATGGAAC
TGATGAATGGGAGCAGTGGTGGAATGCCTTTAATGAGGAAAACCTGTTTTGCTCAGAAGAAATGCCATCTAGTGATGATGAGGCTACTG
CTGACTCTCAACATTCTACTCCTCCAAAAAAGAAGAGAAAGGTAGttGACCCCAAGGACTTTCCTTCAGAATTGCTAAGTTTTTTGAGT
CATGCTGTGTTTAGTAATAGAACTCTTGCTTGCTTTGCTATTTACACCACAAAGGAAAAAGCTGCACTGCTATACAAGAAAATTATGGA
AAAATATTCTGTAACCTTTATAAGTAGGCATAACAGTTATAATCATAACATACTGTTTTTTCTTACTCCACACAGGCATAGAGTGTCTG
CTATTAATAACTATGCTCAAAAATTGTGTACCTTTAGCTTTTTAATTTGTAAAGGGGTTAATAAGGAATATTTGATGTATAGTGCCTTG
ACTAGAGATCATAATCAGCCATACCACATTTGTAGAGGTTTTACTTGCTTTAAAAAACCTCCCACACCTCCCCCTGAACCTGAAACATA
AAATGAATGCAATTGTTGTTGTTAACTTGTTTATTGCAGCTTATAATGGTTACAAATAAAGCAATAGCATCACAAATTTCACAAATAAA
GCATTTTTTTCACTGCATTCTAGTTGTGGTTTGTCCAAACTCATCAATGTATCTTATCATGTCTGGATCCCGTTTAAACTACGCGTAAT
TCAAACAGGGTTCTGGCGTCGTTCTCGTACTGTTTTCCCCAGGCCAGTGCTTTAGCGTTATTGAAAAAGGAAGAGTATGAGTATTCAAC
ATTTCCGTGTCGCCCTTATTCCCTTTTTTGCGGCATTTTGCCTTCCTGTTTTTGCTCACCCAGAAACGCTGGTGAAAGTAAAAGATGCT
GAAGATCAGTTGGGTGCACGAGTGGGTTACATCGAACTGGATCTCAACAGCGGTAAGATCCTTGAGAGTTTTCGCCCCGAAGAACGTTT
TCCAATGATGAGCACTTTTAAAGTTCTGCTATGTGGCGCGGTATTATCCCGTATTGACGCCGGGCAAGAGCAACTCGGTCGCCGCATAC
ACTATTCTCAGAATGACTTGGTTGAGTACTCACCAGTCACAGAAAAGCATCTTACGGATGGCATGACAGTAAGAGAATTATGCAGTGCT
GCCATAACCATGAGTGATAACACTGCGGCCAACTTACTTCTGACAACGATCGGAGGACCGAAGGAGCTAACCGCTTTTTTGCACAACAT
GGGGGATCATGTAACTCGCCTTGATCGTTGGGAACCGGAGCTGAATGAAGCCATACCAAACGACGAGCGTGACACCACGATGCCTGTAG
CAATGGCAACAACGTTGCGCAAACTATTAACTGGCGAACTACTTACTCTAGCTTCCCGGCAACAATTAATAGACTGGATGGAGGCGGAT
AAAGTTGCAGGACCACTTCTGCGCTCGGCCCTTCCGGCTGGCTGGTTTATTGCTGATAAATCTGGAGCCGGTGAGCGTGGGTCTCGCGG
TATCATTGCAGCACTGGGGCCAGATGGTAAGCCCTCCCGTATCGTAGTTATCTACACGACGGGGAGTCAGGCAACTATGGATGAACGAA
ATAGACAGATCGCTGAGATAGGTGCCTCACTGATTAAGCATTGGTAACTGTCAGACCAAGTTTACTCATATATACTTTAGATTGATTTA
AAACTTCATTTTTAATTTAAAAGGATCTAGGTGAAGATCCTTTTTGATAATCTCATGACCAAAATCCCTTAACGTGAGTTTTCGTTCCA
CTGAGCGTCAGACCCCGTAGAAAAGATCAAAGGATCTTCTTGAGATCCTTTTTTTCTGCGCGTAATCTGCTGCTTGCAAACAAAAAAAC
CACCGCTACCAGCGGTGGTTTGTTTGCCGGATCAAGAGCTACCAACTCTTTTTCCGAAGGTAACTGGCTTCAGCAGAGCGCAGATACCA
AATACTGTTCTTCTAGTGTAGCCGTAGTTAGGCCACCACTTCAAGAACTCTGTAGCACCGCCTACATACCTCGCTCTGCTAATCCTGTT
ACCAGTGGCTGCTGCCAGTGGCGATAAGTCGTGTCTTACCGGGTTGGACTCAAGACGATAGTTACCGGATAAGGCGCAGCGGTCGGGCT
GAACGGGGGGTTCGTGCACACAGCCCAGCTTGGAGCGAACGACCTACACCGAACTGAGATACCTACAGCGTGAGCTATGAGAAAGCGCC
ACGCTTCCCGAAGGGAGAAAGGCGGACAGGTATCCGGTAAGCGGCAGGGTCGGAACAGGAGAGCGCACGAGGGAGCTTCCAGGGGGAAA
CGCCTGGTATCTTTATAGTCCTGTCGGGTTTCGCCACCTCTGACTTGAGCGTCGATTTTTGTGATGCTCGTCAGGGGGGCGGAGCCTAT
GGAAAAACGCCAGCAACGCGGCCTTTTTACGGTTCCTGGCCTTTTGCTGGCCTTTTGCTCACATGTTCTTTCCTGCGTTATCCCCTGAT
TCTGTGGATAACCGTATTACCGCCTTTGAGTGAGCTGATACCGCTCGCCGCAGCCGAACGACCGAGCGCAGCGAGTCAGTGAGCGAGGA
AGCGGAAGA

pSL1180-HR-Aag2-loxP-PUb-RFP-loxP-3xFLAG-AGO1

*AGO1* homology with silent mutations at sgRNA cleavage sites

loxP sites

*aae* PUb promoter

RFP

3xFLAG

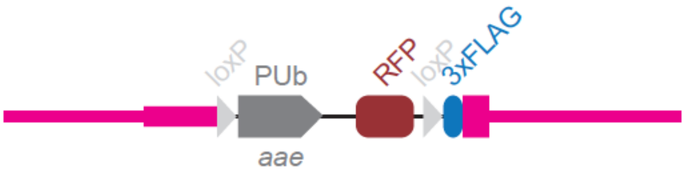


GCGCCCAATACGCAAACCGCCTCTCCCCGCGCGTTGGCCGATTCATTAATGCAGCTGGCACGACAGGTTTCCCGACTGGAAAGCGGGCA
GTGAGCGCAACGCAATTAATGTGAGTTAGCTCACTCATTAGGCACCCCAGGCTTTACACTTTATGCTTCCGGCTCGTATGTTGTGTGGA
ATTGTGAGCGGATAACAATTTCACACAGGAAACAGCTATGACCATGATTACGAATTCGACATGTAGGACATGTGGGGTAGTTCCAGAAA
TCCAGATAAACGATGGACATTTGAATTATCCCCAAACCTCCATCATGTTCTGTTACTTGATATCAACTCAAGGAACCCCGCACTAAAAT
TGAATTTCATGGCTACTATGATAACCGTGTTAAACCGTTTTCTAAAAGTTTCTGTCGCATGACTCGTTACAAATGACGTTACAAATCTA
TAATTTCTATAAATCCCTTCAAAATTGACTCTTGGAAATTTATCTGTATTAAAATTATATAATTATTTTAAAAGCTGGTAAATCACCAA
CATTTTGACATCCATTTCATCCAGAATCGTTCTATTGTTTATGCGGTGAATGCCGAAAAAAATATACCTTCAATGATTCATTGCCGATG
TTTCAACTGAACTCAAACTCAGAAATATGAAATTTCACGCATTTAGATAGATGACTACTCTTTTTTCAGAGTTTTAGTTCACAGCAAAG
ATAAAAGGTAATTCAAATGAGAGCTGCTGTAGAAAGTTTCTCGCTCAATGTATGATCTCGCCTTAAAGCCATTGGTAACCAAGTGTGTA
GCTTGTTGTTACTACGCAAACGAATGGATTTACACGTTACTTAAAAATGCTACAATGAAGATTCTCTATCTCCCAGTGGCGATAGTTAT
ATTTTGGTCCATTCATTTGCTTAAAAACAACGAGCTACACATCAATGGCTTTTAGATCGAGATCACAATTTATGCGAGATTTCCTCAAG
TAATTTATTGCCAAGAAATTTCGTGAATAATAATAATTATAATGATTATTCTGTTTCATGAGCGCATCCATTTTGGAGCGTCCGTAAAA
TGAAACAAAACGGTAAACCGCAACCTGGCTGTTTACTTGCAGCGGTTTTTCGGTACCTAAACTTGCGTACAAGTCGAACTCGTCTAAAA
CATAACGCCTACCCAGGCTGCTGGTAGCGAAACGTTGGGGGGCGATGAAGAAAACTTGTCTCACTCGCCATCATACGAGAGTTTCGCGA
CGAGTTCAGCAGACGTGCGTGTTTAGCGCGACGACGAAAAGTGCAAAAATTCGGCCGTAATTAGTTCGCTTCCGTTTTTCCGAGTGCCT
CCGATCGTTAGCGGACCGTTCCGGGTGTGATTTAAGCGTGAAAATAACTTCGTATAATGTATGCTATACGAAGTTATTATCTTTACATG
TAGCTTGTGCATTGAATCCAATTATAATTTGCCTTGGCACCAGCTGAGCCAGACAAGAAAGAAAGCTTCCCAGAAGTATATCGATTTAG
AAGGGTTGACGTCACTTGCTGACTGCACTAATACAGCAAATGATGCAATTAGAATGATTCAAGTGAAATTCCCAAATTACTGATTTTTC
TCTGGATTTGGTTATCAGATTACATTCGAAGCTAAGATTAGCTACCGAAATTGTCGATCAAATCAGGAAATCCTTTCTCTATCGAAAAA
GGCATTCGCACATCTTCCTCTCTATGCCATATACACGAAGGGTAGGTACATTGACGTCTTTGCCAGAAGTTGAACTGCATCGTTCAAGG
TACAGAATGAACGACTAACAGACACAAGCACGTTTTGCTGTCCATTCAGACACAGGGATGGTACCCATATTCGATCGATATAGAGCCAT
CCAACCGAACAGAGGTATATGTATGAATGTATTGCTGAAATTTTCTAGAAGTACAACCACCACTACGACAGTGTCTATAAAACGCCCCT
GCAAAGGCGAAACCAGCTCAATCGAATACGTTTCCTAGTGGAGTGAACATTACGCGGCCCAAGTAAGCAGTGCCAGTGCAAGTGAAGTG
AAGTCTCTAGTGAAAAAGAGTGATCCAATTAGCCAGAGGAGAAAATTTCAGAGTGAACAAAGCTTTATTCAAAGGACAATTACTATTAA
ATTGGTGAAAGTGCATTTCGGTGAAGGGAATCTTCTAGTGAAGGTAGGTAAATTAATTGATGAAATTATAGCTATGAGCGAAAACTAGT
TTGGTGAATGATTCCTTTGTCTTTGAATGAGCAAACTATTTTCCAAGATGGCGACTATTGAGCTTTGAGTGATTAGTGAAAATTTGCAA
CGCAGTTTCATCATCATTGATAAAACCCAATTGTGATTCACAGCGATAATCATATTTCGTTGAATCATCGCTACTAATTGAATTAAATT
TCTAGAATAATAAGAATAACGTATTTGCTCCGTCACATATCTAAAATAAATATTTTGATGGTTAATTACCCATTAAGGTAATATTAACA
CATATCGAGAAAAACCTTGAGGAAATCGTGAAAACTTGAAGATACGCAATTTCAAAACTACGTAGTTCAAAGTCGAAAACAAGTTAATT
TTTCACTTAAAAGTAGGGCGTTGTTGTGACGTCATCACCTTCAAGTGTATATTTTTCACTTGGCCTGCGACTGCAAACGCAGACAAAGC
AAAACAAGTTTAAAACCTGTCGTGTCGTGCTCGAAGCCAAAGGCAATGAATCAATATCAAATGAGAGTTTGCATTTCACAACCAATTAC
TCAAGCGTTTCCTCGTTTCTTTTTCTGCTCAACAGAGATTTCAACACCAAGCTCGATAAGCTTGTTCGAATCTCGAGTGCGCGCTTCCG
GAGGTATACACCTAGGCGGTACCACTGCAGTGAATTCGGAGCTCTACCGCGGCCGCATGAGCGAGCTGATCAAGGAGAACATGCACATG
AAGCTGTACATGGAGGGCACCGTGAACAACCACCACTTCAAGTGCACATCCGAGGGCGAAGGCAAGCCCTACGAGGGCACCCAGACCAT
GAAGATCAAGGTGGTCGAGGGCGGCCCTCTCCCCTTCGCCTTCGACATCCTGGCTACCAGCTTCATGTACGGCAGCAAAGCCTTCATCA
ACCACACCCAGGGCATCCCCGACTTCTTTAAGCAGTCCTTCCCTGAGGGCTTCACATGGGAGAGAATCACCACATACGAAGACGGGGGC
GTGCTGACCGCTACCCAGGACACCAGCTTCCAGAACGGCTGCATCATCTACAACGTCAAGATCAACGGGGTGAACTTCCCATCCAACGG
CCCTGTGATGCAGAAGAAAACACGCGGCTGGGAGGCCAACACCGAGATGCTGTACCCCGCTGACGGCGGCCTGAGAGGCCACAGCCAGA
TGGCCCTGAAGCTCGTGGGCGGGGGCTACCTGCACTGCTCCTTCAAGACCACATACAGATCCAAGAAACCCGCTAAGAACCTCAAGATG
CCCGGCTTCCACTTCGTGGACCACAGACTGGAAAGAATCAAGGAGGCCGACAAAGAGACCTACGTCGAGCAGCACGAGATGGCTGTGGC
CAAGTACTGCGACCTCCCTAGCAAACTGGGGCACAGATAATTAATTAATAACTTCGTATAATGTATGCTATACGAAGTTATATGGATTA
CAAGGATCACGATGGAGATTACAAGGATCACGATATCGATTACAAGGATGATGATAAGTATCCGGTGGGTCAACGTAAGTACCGTTTCC
GGACGTTTGAAAAAAGTGCAATTCACAAGAGAAGAAAAAAAAAAaGTGTTGgCGAGGGGACCTTCCTTCCTCCCTCTCGTCTCTCTCGCT
GTGTAAATATTTCTTCCCCCATGTGTGTAATTTTCCTGGCCTGCTGCTCTGTTTGGTGCTTCTCTGGACGTTTTTGTGCGTGTGcGTTG
GTGTGATGGGGGGCTTTTCCTGTGTACGGACGAAGAACTTTGTGTCGGTGTGCGTGTGAGATGAGGCCCTCGCCAGAATGAAAATAAAC
CTCATCAACAATGAAAAGTGTCACCACTTTATGGAATAAAAAACGTTTTTATGCAGAAGAGCATGTGAATTATGTATCACTTGGCGATG
GAATTGGGAACATTGATGCTTCCGGACCGATGGAATAATGGAAAACCAGATGTTTCTGCACGTTCTTTGGCACCTTGGCAATGTGTACT
AATCAACACCTCCCTCGAAAAATATACTTGGTGATCAGAATGTTTGTGTTTGTGAATAGTAGGTAGTTTCCGTGTGTGGGTGTGTGTCT
TTGCTAATCATCCTGCTCCAGAAAAAAAGGAAAACCACCAATTTTAATCATCATTTTCTGCGGCTTCGGAGACTGATAAGAGGAATACA
ACTGTTTGCAAACAGTGGATTGAAAACAATGGATAGAAGTTTTATTGGTTCACTTCAGTTCTAAACAGCGGATCCATCTGAAAAGTTAT
ATGAAATTGTACCCCCTTTTCAGTGCTGGGAAATTATTGCCTTTTGTGTCGCTCTTTGCTCACAACTACATACGGAGATGCAGACAGAG
CTCGCATTATGTTGGTGGTGCTCTGTCGCTGGCCATCCGAGAATCGGAAACATCGTGCGGATGGAACATGGAAGAAGAGGACGTTGTAG
GGTTTACCATTATGTATATGCGTTCAGGCGACATCAGACGAGGTGGGTCTTCTCGAAATGATGGGTGGTTGACTGAGTAGCACCTGCTG
TTCATGTGCGGAAATGGAATAACGTGGAGATGCAATGTTCTTCAACGCTAGAAGGATTCGGGGAGACGAAAAGCAAATCCGGTGGCGGC
CGCCTGCAGCTGGCGCCATCGATACGCGTACGTCGCGACCGCGGACATGTACAGAGCTCGAGAAGTACTAGTGGCCACGTGGGCCGTGC

ACCTTAAGCTTGGCACTGGCCGTCGTTTTACAACGTCGTGACTGGGAAAACCCTGGCGTTACCCAACTTAATCGCCTTGCAGCACATCC
CCCTTTCGCCAGCTGGCGTAATAGCGAAGAGGCCCGCACCGATCGCCCTTCCCAACAGTTGCGCAGCCTGAATGGCGAATGGCGCCTGA
TGCGGTATTTTCTCCTTACGCATCTGTGCGGTATTTCACACCGCATACGTCAAAGCAACCATAGTACGCGCCCTGTAGCGGCGCATTAA
GCGCGGCGGGTGTGGTGGTTACGCGCAGCGTGACCGCTACACTTGCCAGCGCCCTAGCGCCCGCTCCTTTCGCTTTCTTCCCTTCCTTT
CTCGCCACGTTCGCCGGCTTTCCCCGTCAAGCTCTAAATCGGGGGCTCCCTTTAGGGTTCCGATTTAGTGCTTTACGGCACCTCGACCC
CAAAAAACTTGATTTGGGTGATGGTTCACGTAGTGGGCCATCGCCCTGATAGACGGTTTTTCGCCCTTTGACGTTGGAGTCCACGTTCT
TTAATAGTGGACTCTTGTTCCAAACTGGAACAACACTCAACCCTATCTCGGGCTATTCTTTTGATTTATAAGGGATTTTGCCGATTTCG
GCCTATTGGTTAAAAAATGAGCTGATTTAACAAAAATTTAACGCGAATTTTAACAAAATATTAACGTTTACAATTTTATGGTGCACTCT
CAGTACAATCTGCTCTGATGCCGCATAGTTAAGCCAGCCCCGACACCCGCCAACACCCGCTGACGCGCCCTGACGGGCTTGTCTGCTCC
CGGCATCCGCTTACAGACAAGCTGTGACCGTCTCCGGGAGCTGCATGTGTCAGAGGTTTTCACCGTCATCACCGAAACGCGCGAGACGA
AAGGGCCTCGTGATACGCCTATTTTTATAGGTTAATGTCATGATAATAATGGTTTCTTAGACGTCAGGTGGCACTTTTCGGGGAAATGT
GCGCGGAACCCCTATTTGTTTATTTTTCTAAATACATTCAAATATGTATCCGCTCATGAGACAATAACCCTGATAAATGCTTCAATAAT
ATTGAAAAAGGAAGAGTATGAGTATTCAACATTTCCGTGTCGCCCTTATTCCCTTTTTTGCGGCATTTTGCCTTCCTGTTTTTGCTCAC
CCAGAAACGCTGGTGAAAGTAAAAGATGCTGAAGATCAGTTGGGTGCACGAGTGGGTTACATCGAACTGGATCTCAACAGCGGTAAGAT
CCTTGAGAGTTTTCGCCCCGAAGAACGTTTTCCAATGATGAGCACTTTTAAAGTTCTGCTATGTGGCGCGGTATTATCCCGTATTGACG
CCGGGCAAGAGCAACTCGGTCGCCGCATACACTATTCTCAGAATGACTTGGTTGAGTACTCACCAGTCACAGAAAAGCATCTTACGGAT
GGCATGACAGTAAGAGAATTATGCAGTGCTGCCATAACCATGAGTGATAACACTGCGGCCAACTTACTTCTGACAACGATCGGAGGACC
GAAGGAGCTAACCGCTTTTTTGCACAACATGGGGGATCATGTAACTCGCCTTGATCGTTGGGAACCGGAGCTGAATGAAGCCATACCAA
ACGACGAGCGTGACACCACGATGCCTGTAGCAATGGCAACAACGTTGCGCAAACTATTAACTGGCGAACTACTTACTCTAGCTTCCCGG
CAACAATTAATAGACTGGATGGAGGCGGATAAAGTTGCAGGACCACTTCTGCGCTCGGCCCTTCCGGCTGGCTGGTTTATTGCTGATAA
ATCTGGAGCCGGTGAGCGTGGGTCTCGCGGTATCATTGCAGCACTGGGGCCAGATGGTAAGCCCTCCCGTATCGTAGTTATCTACACGA
CGGGGAGTCAGGCAACTATGGATGAACGAAATAGACAGATCGCTGAGATAGGTGCCTCACTGATTAAGCATTGGTAACTGTCAGACCAA
GTTTACTCATATATACTTTAGATTGATTTAAAACTTCATTTTTAATTTAAAAGGATCTAGGTGAAGATCCTTTTTGATAATCTCATGAC
CAAAATCCCTTAACGTGAGTTTTCGTTCCACTGAGCGTCAGACCCCGTAGAAAAGATCAAAGGATCTTCTTGAGATCCTTTTTTTCTGC
GCGTAATCTGCTGCTTGCAAACAAAAAAACCACCGCTACCAGCGGTGGTTTGTTTGCCGGATCAAGAGCTACCAACTCTTTTTCCGAAG
GTAACTGGCTTCAGCAGAGCGCAGATACCAAATACTGTTCTTCTAGTGTAGCCGTAGTTAGGCCACCACTTCAAGAACTCTGTAGCACC
GCCTACATACCTCGCTCTGCTAATCCTGTTACCAGTGGCTGCTGCCAGTGGCGATAAGTCGTGTCTTACCGGGTTGGACTCAAGACGAT
AGTTACCGGATAAGGCGCAGCGGTCGGGCTGAACGGGGGGTTCGTGCACACAGCCCAGCTTGGAGCGAACGACCTACACCGAACTGAGA
TACCTACAGCGTGAGCTATGAGAAAGCGCCACGCTTCCCGAAGGGAGAAAGGCGGACAGGTATCCGGTAAGCGGCAGGGTCGGAACAGG
AGAGCGCACGAGGGAGCTTCCAGGGGGAAACGCCTGGTATCTTTATAGTCCTGTCGGGTTTCGCCACCTCTGACTTGAGCGTCGATTTT
TGTGATGCTCGTCAGGGGGGCGGAGCCTATGGAAAAACGCCAGCAACGCGGCCTTTTTACGGTTCCTGGCCTTTTGCTGGCCTTTTGCT
CACATGTTCTTTCCTGCGTTATCCCCTGATTCTGTGGATAACCGTATTACCGCCTTTGAGTGAGCTGATACCGCTCGCCGCAGCCGAAC
GACCGAGCGCAGCGAGTCAGTGAGCGAGGAAGCGGAAGA
